# A Coma Pattern-Based Autofocusing Method Resolves Bacterial Cold Shock Response at Single-Cell Level

**DOI:** 10.1101/2024.10.15.618375

**Authors:** Sihong Li, Zhixin Ma, Yue Yu, Jinjuan Wang, Yaxin Shen, Xiaodong Cui, Xiongfei Fu, Shuqiang Huang

**Affiliations:** State Key Laboratory of Quantitative Synthetic Biology, Shenzhen Institute of Synthetic Biology, Shenzhen Institutes of Advanced Technology, Chinese Academy of Sciences, Shenzhen, 518055, China; University of Chinese Academy of Sciences, Beijing, 100049, China; Department of Physics, The University of Hong Kong, Hong Kong

**Author notes:** These authors contributed equally: Sihong LI, Zhixin MA, Yue YU. Corresponding authors: Shuqiang HUANG.

## Abstract

Imaging-based single-cell physiological profiling holds great potential for uncovering fundamental bacterial cold shock response (CSR) mechanisms, but its application is impeded by severe focus drift during rapid temperature downshifts required for CSR induction. Here, we introduce LUNA (Locking Under Nanoscale Accuracy), an innovative autofocusing method that leverages the coma pattern of detection light to characterize focus drift. LUNA improves the focusing precision down to 3 nm and extends the focusing range to at least 40 times the objective depth-of-focus. These advancements enable us to investigate the complete dynamics of bacterial single-cell CSR, revealing continuous cellular growth and division. We resolve a three-phase adaptation process characterized by distinct growth deceleration dynamics, and show that bacterial cells maintain robust size regulation and coordinate uniform adaptation to cold shock through synchronized growth and elapsed cycles. Notably, a model based on scattering theory reconciles the paradox between the growth lag of batch culture and continuous single-cell growth. These findings fundamentally transform our understanding of bacterial CSR and highlight LUNA’s excellent potential for expanding state-of-the-art research in biology.

## 1. Introduction

The bacterial cold shock response (CSR) is a conserved adaptive mechanism triggered by abrupt temperature downshift^1^. It involves comprehensive physiological regulation that restores cellular growth homeostasis. Investigating CSR is essential for elucidating microbial survival mechanisms across clinical^2^ and industrial settings^3^. This investigation advances fundamental understanding of stress adaptation principles and enables key translational applications, including novel antimicrobial development, enhanced biotechnological protein production, and improved cryopreservation protocols^4^.

Over past decades, CSR has been extensively characterized at the population level^1^. Many bacterial species enter an acclimation phase immediately after cold shock, widely considered a growth arrest, as it is manifested by a constant optical density (OD)^5,6^. This phase is followed by the establishment of a new homeostasis exhibiting a reduced growth rate. While omics approaches have identified key CSR-associated proteins, their multilayered regulation and functional coordination remain poorly understood^1^. Many critical aspects of CSR, such as the physiological significance of the acclimation for bacterial cold adaptation and the specific cellular events occurring to the new homeostasis, remain underexplored. This is largely due to limitations inherent in population-level approaches.

Imaging-based single-cell physiological profiling, featuring high temporal resolution and throughput, has significantly advanced microbiology studies, enabling representative discoveries such as: identification of bacterial persister cell origins^7^, propagation of molecular noise resulting in phenotypic heterogeneity^8^, and validation of the bacterial “adder” behavior (wherein cells add a constant length extension between divisions)^9,10^. However, applying existing single-cell imaging methodologies for unbiased quantitative analysis of CSR faces significant challenges due to specific experimental requirements. Inducing CSR requires rapidly cooling the bacterial culture medium from 37 °C to below 15 °C within minutes. Standard temperature control systems in commercial microscopes lack such rapid cooling capacity. Crucially, attempting this rapid cooling causes severe thermal drift, mechanically displacing the focal plane and resulting in irreversible focus loss.

As with general microscopy imaging workflows, acquiring statistically reliable data in single-cell CSR imaging requires maintaining precise focus across multiple fields of view over hours to days^11^. This necessitates an autofocus system to compensate for inevitable and unpredictable focus drift. A variety of strategies have been developed to this end over the years^12–19^, and can be classified into two groups: imaging-based methods that capture image stacks along the optical axis and assess focus quality metrics to identify the in-focal plane through image analysis, reflection-based approaches that project auxiliary light to sense and measure the resulting reflection changes to directly gauge the distance between the objective and sample (Supplementary Note 1). Nevertheless, current implementations exhibit limited precision and range to accommodate stringent requirements of biological research, particularly the rapid large-span temperature shift required for CSR studies^20^. These limitations similarly hinder other advanced microscopy applications, for instance, imaging across large volume^14^, single-molecule localization^21^, and super-resolution imaging^16^, etc. Meanwhile, computational extended depth-of-field techniques provide axial range extension through single-shot optical encoding or multi-focal fusion, circumventing repetitive autofocusing^22–24^. However, it remains fundamentally inadequate for CSR investigations due to incompatible high-resolution requirements and large drift exceeding its extended depth range.

Herein we propose LUNA (Locking Under Nanoscale Accuracy), a novel reflection-based autofocusing method that breaks through the limitations of the current solutions. LUNA is demonstrated to achieve exceptional focusing precision down to 3 nm while offering an ultra-large focusing range of at least 40 times the depth-of-focus (DOF) of the objective. With the help of LUNA, we investigated the bacterial CSR at the single-cell level by continuously monitoring thousands of individual *Escherichia coli* cells for more than ten hours. Multiple key aspects of bacterial CSR previously obscured in population-level measurements were resolved for the first time. Critically, *E. coli* cells continued to proliferate during the acclimation. Our research revealed significant insights into detailed growth dynamics and facilitating growth mode transition, and further found that they resonate closely with prior molecular-level studies. Remarkably, *E. coli* exhibits a highly uniform, population-wide adaptation to cold shock without employing bet-hedging strategies^25^. These findings substantially advance our understanding of bacterial CSR and provide critical mechanistic insights. Furthermore, we analyzed our batch culture results on the basis of scattering theory, and concluded that the OD lag is actually the outcome of the combined effect of changes in bacterial concentration and volume, which explains the controversies between single-cell and population observations.

## 2. Results

### 2.1. LUNA-based Platform for CSR Single-cell Physiological Profiling

Despite the observed population-level acclimation (growth arrest indicated by OD lag) immediately after cold shock^6^, inherent cellular heterogeneity suggests diverse single-cell behaviors could underlie this net effect (Figure 1a). Several potential scenarios can be proposed: most cells abruptly halt growth simultaneously; some cells might undergo size reduction to adapt to growth-limiting low temperatures, similar to nutrient limitation^10,26^; or conversely, a less sensitive subpopulation may sustain growth, either maintaining or adjusting its growth rate, analogous to antibiotic-tolerant persisters^7^ (Figure 1a). Importantly, these scenarios (individually or in combination) could collectively contribute to the OD lag. The actual dynamics are likely more complex when cell division and viability are also taken into account. Resolving these single-cell behaviors is essential, as it will elucidate the physiological basis of the acclimation and provide key insights into the molecular mechanisms of CSR^1^.

**Figure 1.**
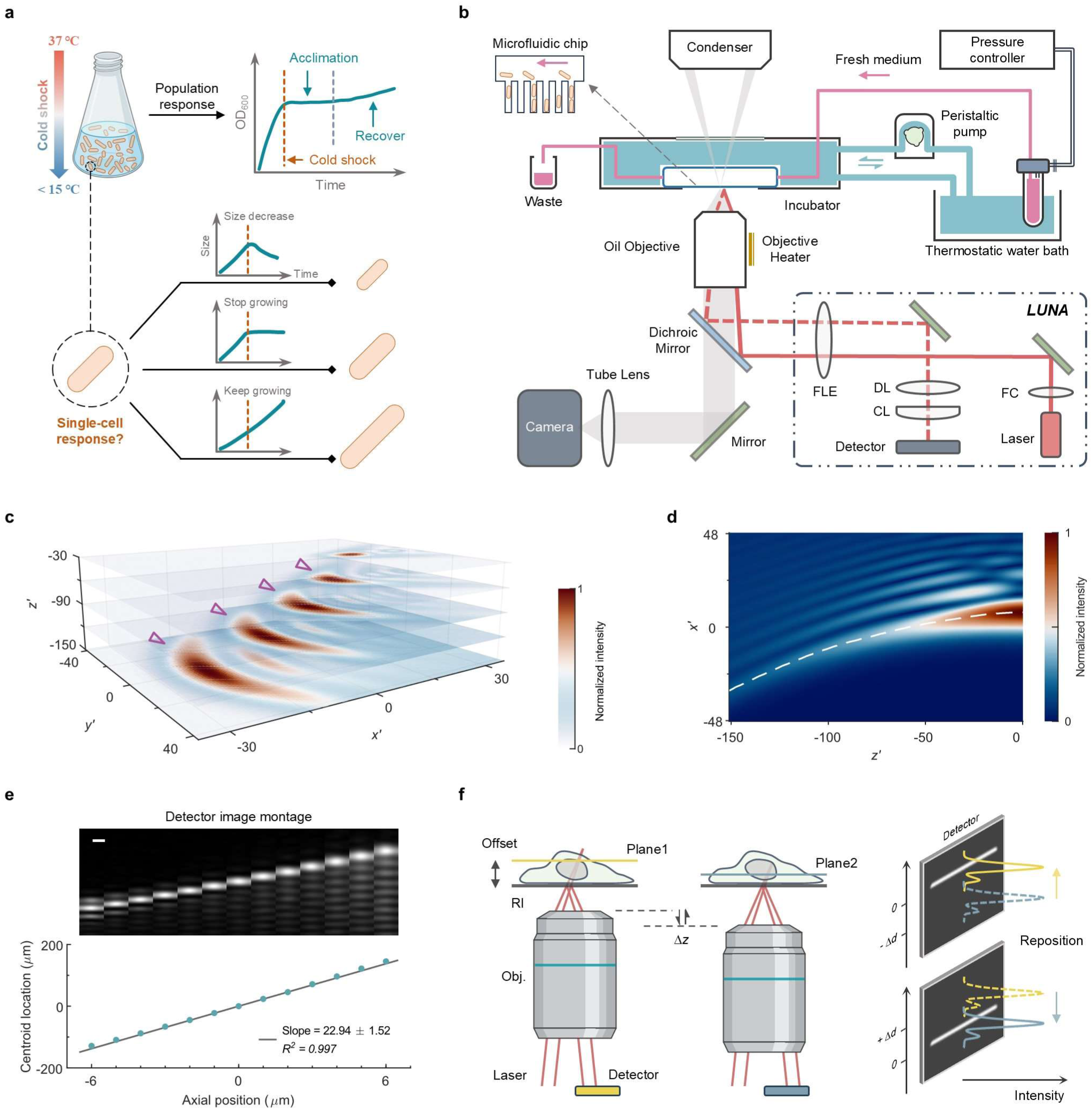
LUNA-based platform for CSR single-cell imaging and illustration of optical principle. **a**, Schematic of hypothesized single-cell bacterial CSR inferred from population-level OD analysis. The red dashed line indicates the time of cold shock (CS) application. After CS, OD ceases to increase, entering an acclimation phase, after which it resumes increase. At the single-cell level, possible cellular size changes following CS include decrease, stasis, or continued increase. **b**, Schematic of our bespoke LUNA-based platform. Diagram includes the phase-contrast imaging unit, microfluidic-compatible temperature control device and LUNA optical path. Circulating water is indicated by the color cyan. In LUNA, laser beam is reflected into the main optical train by a dichroic mirror. Red dashed line: reflected light. FC, fiber collimator; CL, cylindrical lens; DL, detector lens; FLE, focal length extender. The implementation of LUNA follows this design. **c**, Computed intensity distribution in the dimensionless space under 100× objective. Computations were performed using the optical model: An off-axis Gaussian beam propagates through an objective lens of radius *a* and focal length *f*. Coordinates (*x’*, *y’*, *z’*) equals (*kβx, kβy, kβ^2^z*), where *k* is wavenumber, *β* = *a* / *f*. Images at each axial position were individually normalized. Triangle markers indicate the crescent spots. **d**, Computed intensity distribution in x-z space. The dashed line indicates the centroid locations of the brightest crescent spots along the z-axis. Image contrast was adjusted for better exhibition (Gamma = 0.5). **e**, Magnified defocus distance on the detection plane in the ray tracing simulation. Linear regression with slope (95% confidence intervals) and fit goodness is shown. Scale bar, 60 µm. **f**, Working principle of LUNA. Set Plane1 (Plane2) as the interested focal plane by specifying the centroid location of the line spot on the detector as the origin. The defocus distance *Δz* between the target and defocused Plane2 (Plane1) is inferred by the displacement *Δd*. Autofocusing is then achieved by repositioning the spot to the origin. RI, reflection interface; Obj., objective.

To resolve these behaviors, we developed a bespoke LUNA-based platform for single-cell timelapse imaging under rapid temperature change (Methods, Figure 1b). This platform integrates a microfluidic-compatible temperature control device to rapidly and uniformly exert cold shock on individual cells. The microfluidic chip traps single cells in separate microchannels while maintaining steady growth via fresh medium flow^27^, immersed in circulating distilled water for temperature maintenance and change. Precise and rapid cooling is accomplished by replacing the circulating water with ice-cooled water from the thermostatic bath, achieving performance unattainable with commercial air-based thermal controllers (Figure S1, Video S1). To overcome focal drift induced by thermal contraction during rapid temperature shifts, LUNA employs a displacement of coma pattern to monitor focal status and then provide continuous stabilization essential for reliable imaging (Video S2).

### 2.2. LUNA Optical Principle

In conventional reflection-based autofocusing methods, when an objective focuses the focus-aid light beam at the reflection interface, the intensity and contrast of the light spot are the highest. But they decrease rapidly as the defocus distance increases, resulting in a constrained focusing range. Typically, it is also necessary to compensate for the offset distance between the desired focal plane and the reflection interface. However, the additional movement for the compensation leads to more fluctuations that could reduce the focusing accuracy^28^. To alleviate these limitations, for the first time to our knowledge, we introduce coma aberration in the diffraction pattern of the reflected auxiliary laser beam. This allows LUNA to lock onto the focal plan directly rather than introducing the offset, substantially improving focusing accuracy and range.

We investigated a thorough optical model for the propagation of an off-axis laser beam through an ideal lens in the presence of coma aberration^29^ (Methods, Supplementary Note 2). This model gives an analytical expression of light distribution near the focus based on diffraction theory (Figure S2a). The presence of coma aberration causes the intensity distribution of the laser spot to concentrate on one side, forming a high-contrast crescent diffraction pattern (Figure 1c). The centroid locations (center of mass) of the brightest pattern along the z-axis exhibit a monotonic trend over a wide range away from the focal point (Figure 1d), suggesting that the defocus distance can be directly determined by the relative lateral displacement of the centroid location, which can be precisely measured. The contrast of the laser spot away from the focal plane is significantly enhanced compared to aberration-free system with the Airy pattern, thereby improving the accuracy of defocus detection and the focusing range (Figure S2b).

This model serves as a guide for the design and implementation of LUNA (Methods, Figure 1b). To validate and optimize the performance of LUNA, a numerical simulation of the optical setup was carried out based on ray tracing (Methods, Figure S3). The line projection of the crescent spot in the implementation conforms well with the simulation and shows supreme contrast. The simulation shows that LUNA has sufficient sensitivity (Figure 1e, the defocus distance *Δz* can be magnified about 23 times at the detection plane). LUNA allows users to set the focal plane of interest by defining the centroid location on the detector as the origin without the need of setting any offset distance (Figure 1f, Figure S4). This fundamentally decreases the intrinsic motive noises. Additionally, the wide focusing range allows a large tolerance for the offset. In the event of focus drift, a focusing procedure is initiated by repositioning the line spot to the origin through a feedback control.

### 2.3. Performance Analysis of LUNA

The relationship between the measured centroid location and the physical axial position was calibrated (Methods, Figure S5a). LUNA shows an extensive focusing range and good linearity in a particular range (termed as linear range) around the focus, considerably exceeding the corresponding objective DOF. Within the linear range, the lateral displacement *Δd* of the laser spot centroid is given by

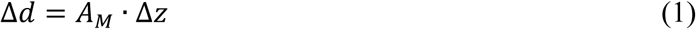

where *A_M_* represents the magnification coefficient. Despite some bias, this relationship can still be used to roughly estimate *Δz* outside the linear range. The measurement repeatability was used to evaluate the degree of consistency during multiple tests. The results highlight the excellent ability of LUNA to accommodate large defocus distances (Table S1), thereby facilitating precise focusing across diverse spatial scales.

A drift threshold value was preset as a constraint criterion to determine the defocus status. LUNA effectively maintains the focal plane within a tight margin, while without compensation the focus drift can escalate to several microns within a dozen minutes (Figure 2a). Our results demonstrate that the focusing accuracy and precision (mean and standard deviation used)^30^ remain small and consistent across different thresholds and objectives (Figure 2b, Figure S5b). Since the feedback autofocusing procedure results in an almost uniform distribution of errors, the measured precision closely approaches the theoretical value (threshold / √3). The best achievable precision is approximately 3 nm in our configuration (Table S1). Apart from the centroid displacement caused by focus drift, other perturbations affecting the precise localization of the centroid primarily arise from temporal fluctuations of the spot intensity. Numerical simulations were conducted to evaluate the effect of perturbations on the centroid location, which is limited to a few nanometers and can be ignored (Methods, Figure S7). Due to the small variance in focusing accuracy, the drift between successive autofocusing procedures is slight (Figure S8). This indicates that LUNA is suitable for common live or time-lapse imaging.

**Figure 2.**
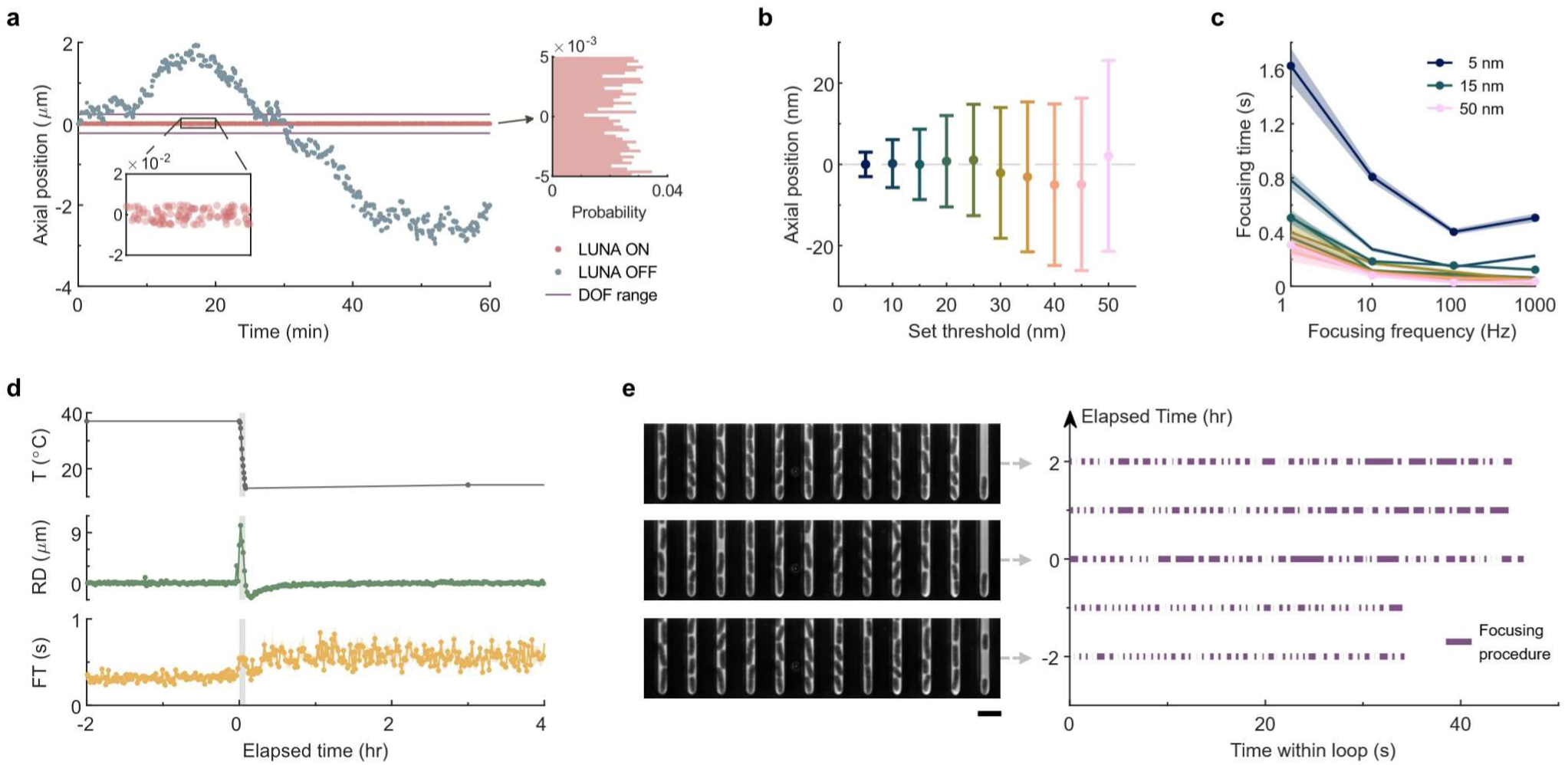
Comprehensive performance metrics of LUNA. **a**, Focus drift monitoring at 1 Hz in the laboratory environment. Right histogram shows the focal plane positions after each autofocusing procedure. **b**, Focusing accuracy and precision assessment. Dots and error bars represent the mean and s.d. values of the focus errors. Color palette in **b**, **c**, different set thresholds. **c**, Focusing time measurement for autofocusing procedures at different intervals. Lines those marked with dots emphasize the representative constraints. **d**, System states over time when imaging bacteria. CS occurs at *t* = 0. T, temperature of cell environment; RD, relative drift of sample stage before autofocusing; FT, corresponding focusing time. Data in scatter form represent the average of all imaged sites (*n* = 45). Shaded region indicates the rapid temperature drop period. Error bands in **c**, **d**, s.e.m. **e**, Representative images and onset of autofocusing procedure. Scale bar, 4 µm. Intervals between consecutive procedures indicate the time consumption of phase-contrast imaging and stage motion.

Short focusing time is another crucial feature of an autofocus system, especially for fast image acquisition. The focusing time was statistically tested under different procedure onset frequencies (Figure 2c, Figure S9). At lower frequencies, the focusing time exhibits a relatively significant variation across different thresholds. This variation can be attributed to an increase in the number of feedbacks needed to stabilize the focus, which arises from the larger axial drift presented. In practical applications, there is a tradeoff between pursuing better precision and shorter time consumption when determining the optimal threshold. Setting a sufficiently large threshold (i.e., greater than 15 nm, which is the optimal value in this work) and a frequency above 10 Hz result in no considerable difference in focusing time under different conditions.

In brief, LUNA locks the focal plane with nanometer-scale precision over an ultra-large range rapidly, ensuring stable focus during long-term imaging for reliable observation of fine subcellular structures and dynamics. These advantages equip us to cope with drastic environmental perturbations during imaging, thereby facilitating the application of LUNA-based platform in bacterial CSR research. Upon initiation of cold shock (CS), the temperature within the sample incubator reached the target of 14 °C from 37 °C within 5 minutes (Figure 2d). This temperature shift caused significant relative drift between consecutive timepoints in the time-lapse imaging (maximum ∼10 μm), much larger than the DOF (Figure 2d, Figure S10). LUNA effectively compensated for this drift, keeping the high-throughput imaging of approximately 1,100 cells simultaneously in focus (Video S2). The average focusing time at each location is around 0.6 seconds, enabling high temporal resolution imaging. (Figure 2d, 2e). Critically, no loss of focus occurred either at the moment of temperature downshift or during subsequent imaging (images in Figure 2e).

### 2.4. Time-resolved Regulation of Growth Without Arrest Under CS

LUNA enables us to investigate the time-resolved behavior of bacterial CSR at the single-cell level. Cell morphological dynamics were extracted from time-lapse image sequences through a self-developed identification algorithm for subsequent statistical analysis (Methods, Supplementary Note 3). Single cells located at the bottom of each microchannel are inherited from one generation to the next after each cell division (Figure 3a). We distinguish the bacterial generations: the one encountering CS (G0), those before CS (GN), and those after CS (G1, G2, etc.). Upon application of CS, sustained division events are still maintained rather than stopping growth as generally believed based on the OD lag (Figure 3a, ∼2 hours in our batch culture). Following acquisition of single-cell data, we addressed how growth and division are affected during CSR by analyzing the growth rate (*λ =* d*L* / d*t* / *L*, where *L* is cell length). Cells exhibit no immediate growth arrest. Constant *λ* before CS confirmed steady growth at 37 °C, while a stable but reduced *λ* after 2 hours indicated a new steady growth at 14 °C (Figure 3b).

**Figure 3.**
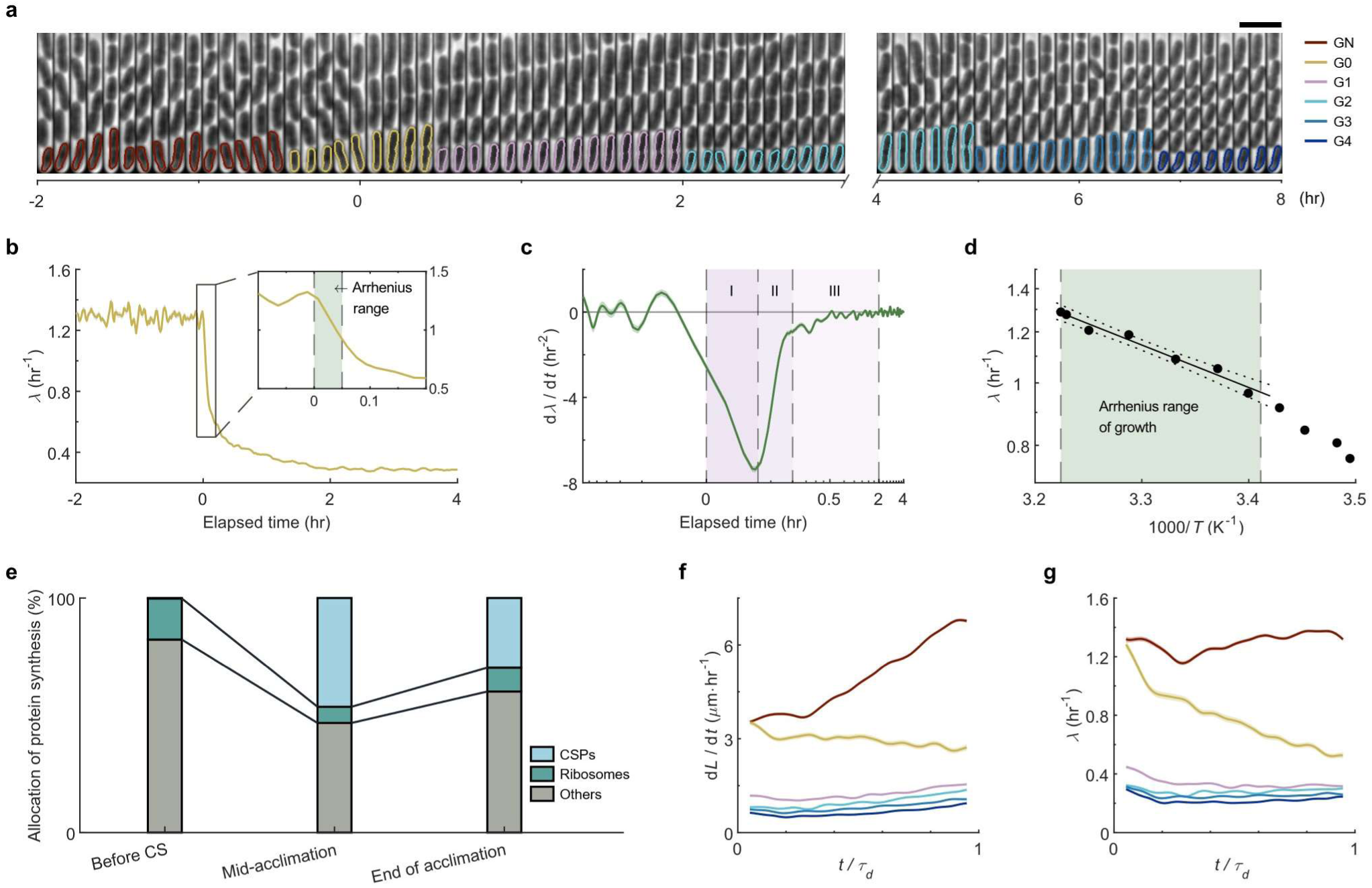
Investigation of single-cell dynamics in CSR using LUNA. **a**, Temporal montage of a single microchannel in the microfluidic chip. CS occurs at *t* = 0. Image was superimposed by cell contours with identified colors according to their generation marks. Timeline unit, hour. Intervals in the left and right panels are 6 and 10 mins, respectively. Scale bar, 4 µm. **b**, Average growth dynamics of cells. Arrhenius range of growth is shaded in inset. **c**, Growth deceleration as function of time. The time window of the three Phases is 0-3 min, 3-10 min and 10-120 min, respectively. Error bands in **b**, **c**, s.e.m. **d**, Temperature dependence of growth rate during the dynamic cooling process. Linear fitting to the reciprocal temperature and logarithm of *λ* is shown as line. Dot lines indicate the 95% prediction bounds. Arrhenius range of growth is shaded. **e**, An illustration of the allocation of different proteins synthesis at different stage during CSR. Percentage data were retrieved from reference^5^. **f**, Average elongation rate across generations as function of normalized interdivision time. **g**, Average growth rate across generations as function of normalized interdivision time. Error bands in **f**, **g**, s.e.m. The color code in **f** and **g** marks the same generation as in **a**.

Analysis of growth deceleration (d*λ* / d*t*) revealed two inflection points at approximately 3- and 10-minute post CS (Figure 3c), delineating three adaptation phases. Phase I (to ∼3 min) involves rapid cooling from 37 ℃ to 20 ℃ (Figure 2d), exhibiting drastic deceleration (approximately -6.25 hr^-2^ on average). Surprisingly, the temperature dependence of *λ* during this dynamic process also follows Arrhenius law observed at the population level in steady-state (Figure 3d), indicating growth deceleration is more likely driven by temperature-dependent reaction kinetics rather than cellular regulated responses^31^. Despite reaching 14 °C within 5 minutes, cells required ∼2 hours to achieve the new steady growth, indicating prolonged physiological adaptation after Phase I. Phase II (to ∼10 min) shows immediate mitigation of growth deceleration (approximately -2.97 hr^-2^ on average), likely mediated by some constitutively expressed cold shock proteins (CSPs, such as CspE) at 37 ℃^32^. These CSPs are generally considered to restore essential biochemical reactions during the acclimation^33^. Sustained elongation of already initiated protein synthesis before CS^34.35^ may also contribute. Phase III continues until the end of the acclimation (to ∼120 min), during which the temperature is kept constant at 14 ℃. The further mitigated deceleration (approximately -0.18 hr^-2^ on average) suggesting more complex physiological regulation. Reanalysis of the protein synthesis dynamics after CS^5^ revealed increase in CSPs synthesis (Figure 3e). These CSPs can recover the protein translation more efficiently^36,37^, likely contributing to the mitigation of growth deceleration during Phase III. Following CS, ribosomes become underutilized as translation initiation is significantly impaired at low temperature^5^. Although ribosome synthesis is reduced during this period, we speculate that this reduction does not exacerbate the growth deceleration, as the dominant mitigating effect is attributed to the upregulated CSPs. Towards the end of acclimation, a new balance between CSPs and ribosomes is established, enabling establishment of new homeostasis (Figure 3b).

Growth mode analysis corroborated our physiological regulatory speculations in Phase III (Figure 3f, 3g). *E. coli* cells sustain exponential growth during steady-state^10^, which is based on the view of “autocatalytic loop” of biomass synthesis^38^. This means that a fixed portion of total newly synthesized proteins is allocated to the ribosomes^39^. The progressive decline in new ribosome synthesis after CS prevents the “autocatalytic loop”, manifested by immediate deviation from exponential growth (Figure 3e). As the synthesis recovers in late phase III, a new homeostasis of “autocatalytic loop” in biomass synthesis is achieved, i.e., the cell completes cold adaptation and resumes exponential growth.

### 2.5. Coordinated and Robust Single-cell Adaptation to CS

The observed disruptions to exponential growth of individual cells suggest that the shock timing within cell-cycle may be determinant; we therefore assessed whether the specific cell-cycle stage at the time of shock influences subsequent heterogeneity and synchrony. To resolve sub-cell-cycle responses in detail, we grouped cell lineages by the relative CS onset time since cell birth of G0, quantitatively defined as *ξ* (Figure 4a). This *ξ*-based grouping revealed the immediate transitions to linear growth in growth mode (Figure S12), likely driven by CS-induced reprogramming of ribosome allocation (Figure 3e). Such transitions may consequently perturb cell size homeostasis, prompting us to investigate its coordination with size regulation mechanisms^40^.

**Figure 4.**
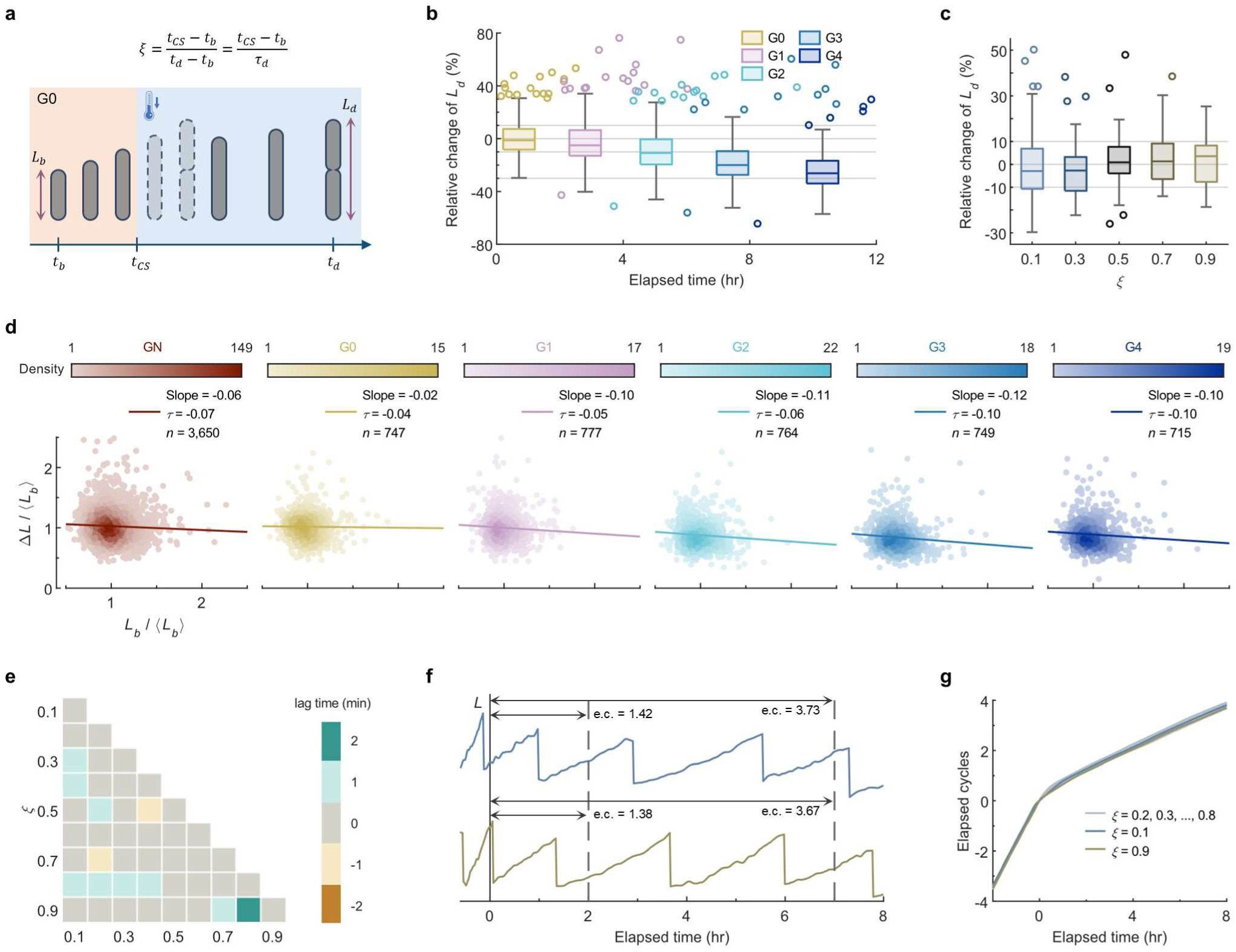
Cell size regulation and growth synchronism under CS. **a**, Notations in cell cycle. *ξ*, relative CS onset time since cell birth; *τ*_d_, interdivision time. Dashed cells, expected exponential growth in the steady-state. **b**, Percentage change of division length of each generation after CS compares to GN. Data are presented as box plots with the edges at the 0.25 and 0.75 quantiles, the central line at the median, and the whiskers at the minima and maxima within 1.5× interquartile range from the box. Circles, outliers outside the whiskers. Each box is located at the average division time of that generation. **c**, Percentage change of division length of each group between G0 and their corresponding predecessors. Data are presented as box plots as in **b**. **d**, Relationship between length extension and birth length across generations (rescaled by the expectation of *L_b_*). Gradient colors indicate data density of each generation. Linear regressions of all the measured cycles and Kendall’s *τ* are shown. *n* represents cycle number. **e**, Lag matrix for distinct groups of *ξ*. Values were computed by locating the cross-correlation maximum. **f**, Representative lineages with synchronized elapsed cycles (*ξ* is 0.10 and 0.90, respectively). e.c., elapsed cycles. **g**, Elapsed cycle numbers of all the grouped lineages as functions of time. Errors shown in Figure S14b.

Bacterial cell size is a sensitive indicator of environmental stress^20,41–44^. We observed that they try to maintain sufficient size (length) extensions in G0 as in the normal steady-state (Figure 4b, Table S2). The division length (*L_d_*) difference between G0 and its corresponding predecessor of each cell is small (median of -1.3%). Furthermore, *L_d_* in G0 remains invariant across lineages with different *ξ* (Figure 4c), consistent with the ungrouped data. The cells then take an additional 3-4 generations for length stabilization to a reduced value (Figure 4b). Besides, *E. coli* cells maintain a narrowly distributed size under the steady-state as the “adder” behavior^9,10,45^, which means length extension Δ*L* within a generation is independent of length at birth *L_b_*. Kendall’s *τ* between rescaled Δ*L* and *L_b_* and the linear regression were examined, showing almost no significant correlation (Figure 4d, |*τ*| ≤ 0.1 for all generations). The size distribution lacks distinct subpopulations, and the standard deviation is almost constant across generations, indicating no substantial increase in cell size variability (Figure 4d, Table S2). These data suggest that this “adder” behavior could be maintained consistently during CSR, indicating that the intrinsic size regulation mechanisms continue to function robustly, thus ensuring proper adaptation of cell size to the stress.

Further studies on cells with distinct *ξ* and their progeny after CS revealed that cells exhibit highly similar behaviors. The changes in *λ* are highly synchronized in time, with the exact time lag between any pair of *ξ* not exceeding 2 min (Figure 4e, Figure S13). Alongside consistent size variation, *E. coli* exhibits no stress-specific subpopulations^7^ (e.g., persisters), or CS-induced cell death (Video S3) indicating a sacrificial subpopulation^46^. This absence of bet-hedging strategies suggests that adaptation to CS is instead achieved through a collective and synchronized response. Surprisingly, the exact number of elapsed cycles at any moment since the CS event is also consistent across different cells (Figure 4f, 4g, Figure S14), implying the division timing is dependent of *ξ*. We verified the consistency of the elapsed cycle deviation over time. This alignment is reflected in the narrow distribution of elapsed cycles for a given time point, which persists over successive generations. Compared with the steady-state, the deviation after CS does not increase significantly and is still within a small range (Figure S14), indicating that cells actively maintain alignment in cycle progression. In other words, cells encountering a relatively late CSR will “accelerate” divisions to maintain progression of division cycle (Figure 4f).

### 2.6. Scattering Theory and Modeling Explanation of Proliferation Persistence During OD Lag

The discoveries from single-cell experiments revealed that *E. coli* continues to proliferate after CS, prompting us to reexamine batch culture experiments (Methods). In our results, the measured OD_600_ curve indeed showed an acclimation immediately following the rapid temperature downshifting (Figure 5a, Figure S16a). Nevertheless, the cell concentration (*C*) continued to increase at a lower rate during the acclimation. Meanwhile, the cell volume (*V*) decreases. Consistent with single-cell experiments, these results confirmed that bacterial divisions do not cease after CS – contrary to conventional understanding.

**Figure 5.**
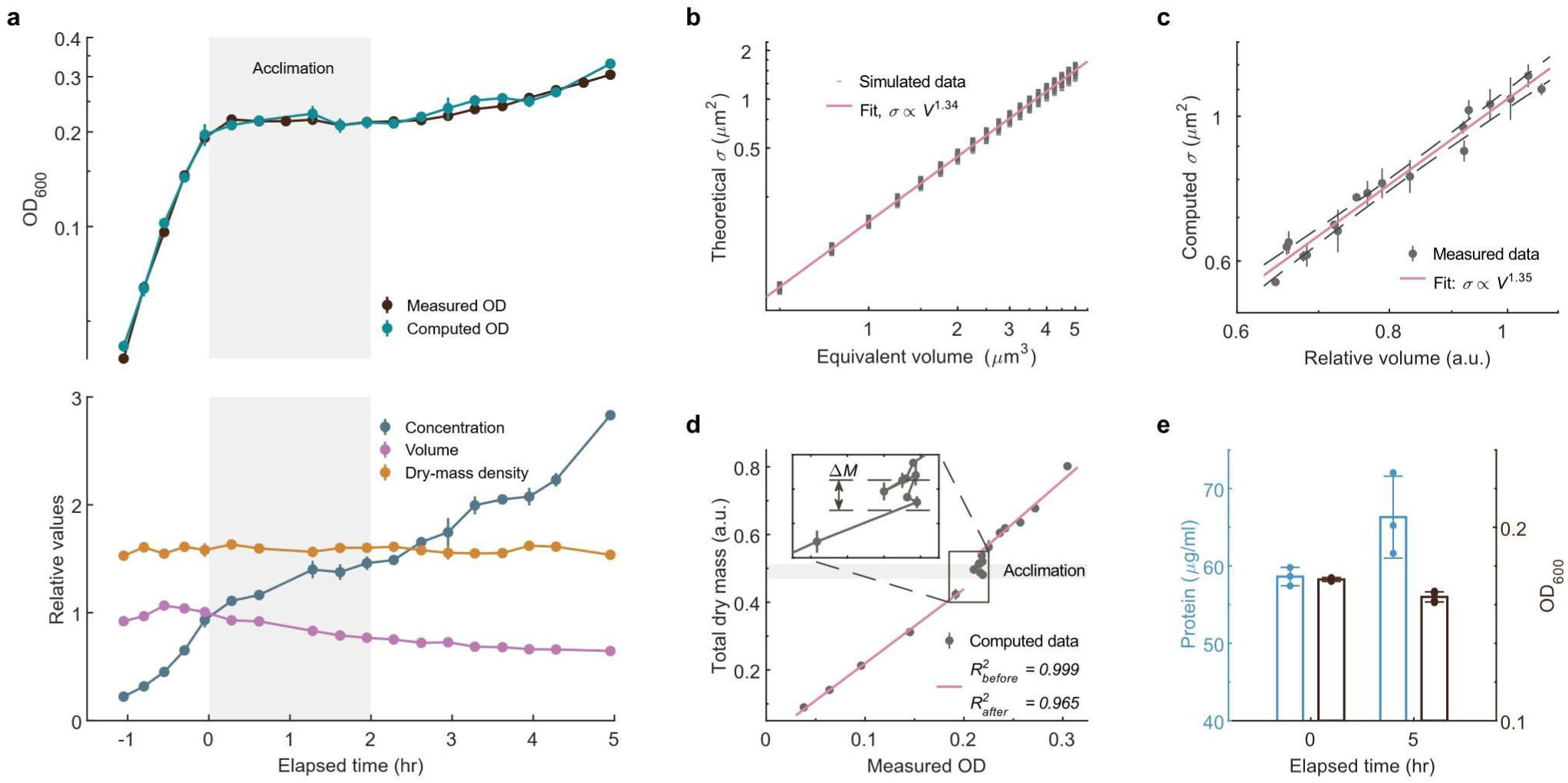
Analysis of CS experiments of batch culture. **a**, Measurements with temperature dropped to 14 ℃. Measured and computed OD_600_ values (top). Computed data was scaled at *t0*. Bottom: cell concentrations and average volumes extracted from the flow cytometer, normalized to *t0* and average value before CS, respectively. Relative cell density *ρ_dry_* was computed and scaled for present. Shaded region indicates the acclimation. **b**, Theoretical *σ* across a range of equivalent cell volumes of simulated cells. Power exponent was determined by linear regression on the logarithmically transformed data (*R*^2^ = 0.996, confidence interval is ±0.01). **c**, Computed *σ* from experimental data as function of relative volume. Power exponent was obtained as same process in **b** (*R*^2^ = 0.971, confidence interval is ±0.12). Dashed lines indicate the 95% prediction bounds. a.u., arbitrary units. **d**, Computed total dry mass plotted as a function of OD. The solid line represents a linear fit forced through the origin, based on the physical constraint that mass must be zero at zero OD. Fitting for the acclimation has a *R*^2^ of -0.263 (Pearson correlation coefficient -0.00). Mass increment during the acclimation is indicated as the height of the shaded area and marked in the inset. Error bars in **a**, **c, d**, s.d. **e**, Total protein measurement of MG1655 strain. Temperature drops to 12 ℃ were performed.

Single-cell experiments show that CS doesn’t cause significant cell death (Video S3). This means OD lag is not a combined effect of partial cell lysis and partial growth. Therefore, a complex question is raised: given that OD is a widely accepted indicator of total dry mass in experiments^47,48^, why does the bacterium’s growth continue while the OD levels off? In theory, OD value involves not only the bacterial concentration, but also light scattering by individual cells. Rayleigh scattering is the predominant scattering of light by small particles compared to the wavelength of incident light. The Rayleigh-Gans approximation can provide more accurate results if the interference scattered from different parts of the scatterer is taken into account^49^ (Methods, Supplementary Note 4, Figure S15). Numerical simulations were performed to compute the total scattering cross section *σ* of well-dispersed typical *E. coli* cells. The results show that *σ* is not correlated to the aspect ratio (Pearson correlation coefficient -0.07) but is theoretically proportional to *V*^1.34^ (Figure 5b, Figure S15c). OD value is then given by *KσC*, where *K* is constant. A concise dependency can be extracted as:

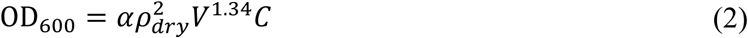

where *α* is constant coefficient depending on the measurement conditions, *ρ_dry_* is the cellular dry-mass density.

In the batch culture, *σ* could be computed from the ratio of OD to *C*. It has a power relationship with *V*, which agrees well with the theoretical relationship (Figure 5c, Figure S16b). This also indicates that *ρ_dry_* remains stable all the time even when cells experiencing CS^38^ (a relative value for *ρ_dry_* is computed from equation (2) by taking the square root of the ratio of the measured OD_600_ to the product of the measured *V*^1.34^ and *C*). Therefore, a theoretical OD curve can be deduced according to equation (2) (Figure 5a, Figure S16a). The relationship holds when single scattering is applicable, which corresponds to the volume concentration of scatterer is less than 0.001^50^ (equivalent to OD_600_ < 0.5 for *E. coli*). Batch culture studies without CS verified that when *C* is high, multiple scattering must be considered, and the cells become physiologically different, leading to increased deviations (Figure S17)^26,51^.

To further verify the validity of equation (2), total dry mass per ml (*M* = *ρ_dry_VC*) as a function of measured OD is also computed. In a steady-state, *M* shows an excellent linear relationship with OD (Figure 5d, Figure S16c, 17b). When bacterial population reaches a new steady-state at a lower temperature, the linear relationship can be well-fitted again. Some mass increment (Δ*M*) during the acclimation indicates population growth. In contrast, the corresponding OD may even decrease slightly due to the complicated interplay (Figure 5e, Figure S18). A protein quantification measurement verified that total protein during the acclimation still increased by about 13%, consistent with the trend and value of computed Δ*M* (Methods). Overall, these results suggest that OD is not an appropriate indicator of total dry mass under perturbed growing environment or unsteady growth, particularly for bacterial CS.

## 3. Discussion

Environmental stress profoundly influences bacterial physiology, population dynamics, and microbial evolution. A key example is CS, a sudden temperature downshift known to induce growth arrest and upregulate responsive genes in *E. coli*. High spatiotemporal imaging is the pivot for characterizing single-cell physiological dynamics, where maintaining the focus precisely during environmental fluctuations remains challenging (Video S1). Herein, we developed an integrated platform for uninterrupted high-throughput time-lapse imaging of bacterial cells at 1-minute intervals. Under the instruction of an optical model with coma aberration, we developed the LUNA method with nanoscale precision and ultra-large focusing range, which can lock the focus directly and rapidly. Four representative objectives were tested to evaluate LUNA’s comprehensive performance metrics based on a range of criteria^52,53^. In our microscope, the precision is limited by the accuracy of the servo motor used, which can be further improved by employing piezoelectric motors with higher positioning accuracy to drive axial movement. The in-depth time consumption analysis demonstrated the applicability of LUNA in tracking defocus variations either in real-time or time-lapse imaging. Although the focusing time is sufficient for our experiments, it can be reduced even further with faster motors and well-calibrated feedback control algorithms.

LUNA is successfully exploited in more extreme out-of-focus scenarios caused by severe temperature fluctuations to study bacterial CSR at the single-cell level. The physiological profiling was performed and revealed many characteristics of CSR that have been covered by population-level OD measurement. A continuous three-phase growth decrement phenomenon in acclimation aligns with molecular-level mechanisms, with cells sustaining both growth and division throughout. Each phase displayed a distinct growth deceleration pattern: Phase I is primarily governed by Arrhenius law kinetics. Subsequently, CSR is reflected in single-cell growth dynamics by the significant mitigation of the reduction in *λ* when the temperature drops below 20 ℃ (widely regarded as a triggering threshold of CSR, and marking Phase II entry), with both Phase II and III explained by the dynamics of protein synthesis and the regulatory roles of known CSPs. Effects related to ribosome allocation may also explain the transition to a linear growth mode. Another noteworthy finding is that despite attenuated growth, bacterial cells maintain size homeostasis and synchronize cell cycle progression, with no evidence of subpopulation formation or reduced cellular viability. This homogeneous adaptive response suggests that *E. coli* employs a collective stress-resistant strategy to tolerate transient temperature stress. Specifically, this strategy involves the activation of a repertoire of pre-synthesized sensory and responsive molecules that persist in a standby state^32,54^ to counteract abrupt temperature fluctuations, thereby contributing to the ecological resilience and community stability of bacterial populations^55^. In addition, our results show that OD is not an accurate growth indicator in batch culture studies when the cell phenotype (size and division) is not maintained. We summarized the OD value dependence and explained the paradoxical phenomenon of growth arrest observed in population. Another scenario where OD may decrease occurs during nutrient downshift, in which the bacteria size reduces^56^.

From the perspective of evolution, it is reasonable for bacteria to keep growing and dividing after CS. Individuals who can increase the population size during the acclimation can gain a competitive advantage when entering a new steady-state. Based on the retention of some CSPs and nucleic acids at optimal growth temperatures, it is foreseeable that bacteria may have developed the ability to sustain growth after CS long time ago. Furthermore, such adaptive strategies can also cope with transient and periodic temperature fluctuations, allowing cells to recover more swiftly to optimal growth following brief CS events without the need to reboot their physiological activities. To further understand the mechanisms underneath those phenomena, more physiological changes involving molecular mechanism exploration can be investigated^57,58^. Looking forward, numerous questions regarding CSR remain unresolved. For instance, why do distinct bacterial species employ divergent adaptive strategies during CSR^1^? How do bacterial cells modulate their physiological states upon the reversion of temperature to optimal growth conditions? Leveraging the LUNA-based platform, single-cell investigations hold the potential to provide critical insights, facilitating a more systematic and rigorous exploration of these unresolved issues.

Beyond the instance in this work, LUNA offers excellent precision and can overcome the limitations of existing autofocusing solutions, thus enhancing many cutting-edge microscopy technologies, including super-resolution, single-molecule imaging, etc. Given its large focusing range, LUNA is capable of easily locking the focus in microscopy imaging across large volumes or at greater depths, such as multiphoton or light-sheet microscopy. The implementation of our method avoids complicated optics, alignment, and operation routines, posing its universal applicability to general microscopy techniques. As a result, we anticipate that LUNA will prevail in various research and industrial applications.

## Methods

### Optical setup of LUNA-based platform

The optomechanical design was performed using CAD software (Dassault Systemes, SolidWorks). An infinity imaging system based on a commercialized microscope frame (Applied Scientific Instrumentation, RAMM) and a set of precision machined components was custom-built. The specimen was mounted on a motorized stage (MS2000) for planar motion. Axial movement of the objective was achieved by a motorized actuator with an encoder resolution of 5.5 nm (LS50). An anti-backlash routine was introduced at each small move to advantage for precision movement. Kohler illumination with a neutral white light source (Thorlabs, MNWHL4) was designed to provide brightfield and phase-contrast imaging mode. High-resolution images were collected using a sCMOS camera (PCO, Panda 4.2) with a 6.5 µm pixel size.

LUNA was installed in this imaging system. The light source was a fiber-coupled laser with a wavelength of 650 nm (output power < 5 mw), and was collimated to parallel beam by a fiber collimator. The wavelength was chosen to avoid crosstalk with the illumination for our bacteria imaging experiments and for a lower cost, although an infrared wavelength for more general applications could also be selected. An aspheric lens was inserted as the focal length extender (FLE), shifting the focal point of the laser beam to optimize the focusing range. The off-axis laser beam was introduced into the main optical train by a shortpass dichroic mirror (Semrock, FF625-SDi01). The diffraction pattern at the reflection plane was projected on a detector (Thorlabs, DCC1545M), whose pixel size is 5.2 µm. A cylindrical lens was placed before the detector to shape the diffraction pattern one-dimensionally to improve the detection accuracy. The full width at half maximum of the line projection of the crescent spot was approximately 30 µm wide, ensuring adequate subpixel accuracy. During the autofocusing procedure, a practical proportional-integral controller was used. The exposure time of the detector was set to 3∼5 ms to avoid overexposure. Although LUNA was implemented on a bespoke imaging system, it is also compatible with other commercial microscopes with infinity optics. Four objective lenses were tested for LUNA’s performance (Table S1). Unless otherwise stated, data presented in the text was collected from the 100× oil objective.

The microfluidic chip was fixed into a custom-designed incubator, which adapts to the motorized stage. The incubator can be filled with distilled water in which the microfluidic chip is submerged to maximum heat conduction. Rubber rings are used in the design of the incubator to prevent water seepage. The sealable incubator has two pairs of fluidic conduits: fresh medium flows through the chip for cell growth by a pressure controller (FluidicLab, PC1); distilled water is circulated into the incubator for temperature change and maintenance. The culture temperature was maintained by the circulation of distilled water in an external thermostatic water bath driven by a peristaltic pump. The temperature of water inside the incubator was set to target during the experiments. In addition, the imaging objective was equipped with a lens heater.

### Numerical computations of the optical model

Diffraction patterns near the focus of the optical system with circular aperture and coma aberrations can be handled by analyzing the wavefront distortion of an off-axis beam (Supplementary Note 2). The objective lens is treated as an ideal lens for the model demonstration. Consider an incident collimated Gaussian beam passing through the lens of radius *a* and focal length *f* (Figure S2a). According to the Huygens-Fresnel principle, the light disturbance at an arbitrary observation point *P* in the vicinity of the focal point is determined by summing the contribution of each point *Q* located where the incident Gaussian beam intersects the reference sphere on the pupil plane^29^. The coma aberration function can also be expressed in the polar coordinates (*ρ*, *θ*) whose origin is the center of the pupil by *Φ*(*ρ*, *θ*) = *A_C_ρ*^3^ cos*θ*, where *A_C_* is the coefficient of primary coma^59^. The complex amplitude *U*(*P*) is determined by calculating the diffraction integral over the intersection area *S* on the reference sphere^29^:

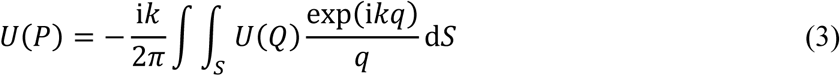

where *k* is the wavenumber, and *q* is the distance between points *Q* and *P*. For the incident beam with small off-axis amount involved, the inclination factor is approximated as unity and therefore is omitted in equation (3). The polar coordinates (*ρ*, *θ*) are translated to (*r*, *φ*) with the origin of which (*ρ_0_*, *θ_0_*) is situated at the beam center for convenience. The amplitude distribution of the Gaussian beam is given by *A_G_* exp(-*a*^2^*R*^2^ / *ω*^2^), where *A_G_* is constant amplitude, and *ω* is beam waist. Furthermore, cylindrical coordinates (*u*, *v*, *ψ*) are introduced to specify an arbitrary point *P* in the image field where *ψ* is the azimuth at the focal plane (*x, y*), the dimensionless coordinates *v* = *ka* (*x*^2^ + *y*^2^)^1/2^ / *f* = *kβ* (*x*^2^ + *y*^2^)^1/2^ and *u* = *kβ^2^z*. Then *U*(*P*) is given by:

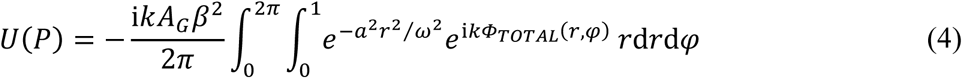

The total phase term of *Φ_TOTAL_* is defined by combining the like terms:

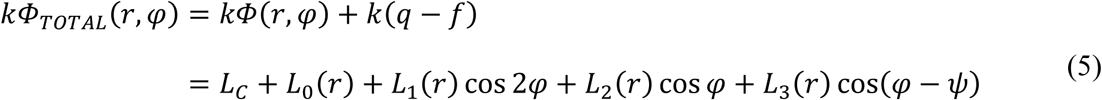

Define *η* = (*L_2_*^2^ + *L*^2^ + 2*L_2_L_3_* cos*ψ*)^1/2^, tan*χ* = -*L_3_* sin*ψ* / (*L_2_* + *L_3_* cos*ψ*). The integral difficulty of the intensity is simplified by expanding the exponential term exp(i*kΦ_TOTAL_*) with the help of the Jacobi-Anger identity and Graf’s addition theorem, substituting the expended series in the integration, and integrating term-wise against *φ*. The intensity distribution of the diffraction pattern is developed in an infinite series:

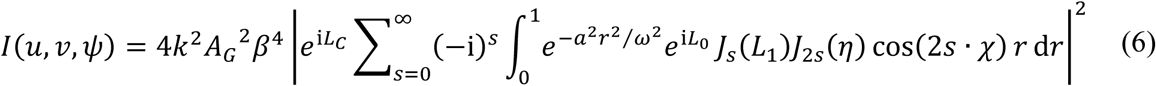

where *J* is a Bessel function of first kind. The term *s* = 0 is taken with the factor 1/2. Tedious calculation can be avoided because four terms in this expansion would suffice to keep the intensity variation within a few percent. The simplification greatly speeds up the numerical calculation of the intensity distribution. In the numerical computation, the selection of each parameter is as close to the actual optical path configuration as possible, such as *β* = 1.25 for the 100× objective lens, relative beam position (*ρ_0_* = 0.1, *θ_0_* = 0°), and coma coefficient *A_C_* = 16*π* / *k*. The coefficient *A_C_* is proportional to the tilt angle of the beam incident on the objective, and we have chosen a more representative value here.

### Ray tracing simulation of the model

The optical design and numerical simulation of LUNA’s optics were carried out based on ray tracing software (Zemax, OpticStudio), to validate LUNA’s robustness under real-world optical limitations. A set of lenses was used to combine to substitute objective with an equivalent magnification of 100×. The other parameters of the optical elements were set exactly the same as the optical setup. For off-axis incidence, the normalized radial coordinate of the beam center at the objective rear aperture was specified to be approximately 0.15 (Figure S3b). To ensure full beam capture by the substitute lens system, the input beam size was reduced at the rear aperture via slight positional adjustment. A reflection surface was placed at the imaging space of the objective. Sequential intensity distributions on the reflection and detection plane were recorded as the reflection surface changes along the optical axis.

### Calibration of defocus dependence

The measurement was carried out by scanning the objective from bottom to top along the optical axis around the focal plane. The reflected images on the detector were repeatedly recorded five times in repetitions at each step. The centroid coordinates of the light spot at sub-pixel precision were then computed from the captured images through basic morphological processing. The linear range was determined by finding the maximum *N* multiples of DOF while maintaining the fitting root mean squared error (*RMSE*) less than 1 (Figure S6). The strict criteria for the linear range determination were selected to obtain high focusing sensitivity and accuracy near the focal plane.

Measurement repeatability is defined as the proportion of the total variation accounted for by the system variation to assess the system’s measurement variation^60^. It indicates the degree of consistency during multiple measurements. The pooled standard deviation was used to calculate this variation underlying the monotonic response curve. A ratio below 10% is indicative of satisfactory system performance^61^. All four test objectives demonstrate measurement repeatability of <2%, confirming the robustness of the calibration relationship.

### Performance test of LUNA

Timelapse tests with specific time intervals (1 second) were carried out and lasted for one hour for the four objectives. Discrete drift thresholds varying from 5 nm to 50 nm at intervals of 5 nm were tested in each condition. In tests evaluating the focusing time required by each autofocusing procedure, the time interval (i.e., the reciprocal of the focusing frequency) was altered. The objective and sample temperature were maintained at 37 ℃.

The extent of the perturbations can be reflected in the temporal fluctuations of spot intensity on the same focal plane. This fluctuation may originate from laser instability, thermal effects in optical elements, etc. The distribution parameters were extracted from the detector images. Random Gaussian noise conforming to this distribution was generated and superimposed onto different spot images to simulate the fluctuations, and the simulation was repeated 100 times on each image. Errors of the centroid locations were employed to evaluate the impact of perturbations.

### Single-cell culture on microfluidic device

A microfluidic chip was employed to track the long-term growth of bacteria, the structure of which follows that of previously published: it consists of a series of microchannels perpendicular to the main trench that perfuse the fresh medium^62^. The chip was fabricated based on soft lithography technique by mixing dimethylsiloxane monomer and curing agent (Dow Corning, SYLGARD 184 Silicone Elastomer Kit) in a ratio of 10:1, and then pouring into a mould, degassing and curing at 80 °C for one hour. The PDMS chip was punched and bonded to a clean coverslip by air plasma treatment (Harrick Plasma, PDC-002) for 30 seconds, followed by baking at 80 ℃ for five minutes. a) *E. coli* K12 strains MC4100 and MG1655 were measured (MC4100 is presented in the text), in which the flagella structural protein gene *fliC* and fimbriae structural protein gene *fimA* of MG1655 were knocked out to prevent their escape from the microchannels. EZ rich defined medium (RDM) supplement with 0.4% glucose was used^63^. The overnight culture grown from several colonies was subcultured twice and centrifuged at a sufficiently high density. The concentrated culture was injected into the cleaned chip passivated with bovine serum albumin, and the chip was spun to load cells into the microchannels. Fresh medium was infused into the chip at a rate of 500 µL/hr by the pressure controller equipped with a 0.22 μm filter. Next, the incubator with the chip was sealed and placed on the motorized stage for time-lapse imaging.

### Time-lapse imaging of bacterial under CS

We developed a control software using MATLAB for the time-lapse imaging in the bespoke platform. Users could set up imaging conditions and perform automation of experiments. The autofocusing control is integrated, with the user only needing to set the desired plane as the focal plane. The output laser power of LUNA at the objective front aperture was adjusted to ∼5 µw, and the threshold was set as 50 nm.

Phase-contrast images were acquired with the 100× objective. Multiple microchannel sites (around 45 in each repeat experiment) were imaged every minute for 24-30 hours. Each image acquisition started right after an autofocusing procedure. Approximately 1,100 single-cell data in total were collected in each repeat. Cells were cultured at 37 ℃ to steady-state for about four hours before CS. By mixing ice into the external water bath, the incubator temperature can be downshifted to 14 ℃ within 5 minutes and maintained until the end of the time-lapse experiment. Although the exact temperature around bacterial cells in the microfluidic chip could not be directly measured, the high thermal conductivity at the water-glass interface ensures rapid thermal equilibration. When the temperature was lowered, the objective lens heater was switched off, causing the objective to approach ambient temperature while the sample was maintained at the target low temperature; this temperature mismatch resulted in a slightly longer focusing time (Figure 2d). At least 3 repeated experiments were carried out for each of the two strains.

### Image analysis and data process

Hundreds of cells at the bottom of microchannels were imaged sequentially. We developed a custom algorithm routine to analyze phase-contrast images using MATLAB: Microfluidic Architecture based Resilient Contour Expansion Analysis (MARCEA) (Figure S11a). Raw images were denoised and segmented to identify individual cells. The contours of the segmented cells were iteratively adjusted based on a rod-shaped model to adapt to the deformation of the cells, thereby extracting their accurate spatial features. Microchannels were automatically aligned for cell tracking (Figure S11b). A cell division event was determined according to the large drop of the width and intensity at the split position. Using the processing routine, complete analysis of ∼65,000 images costs ∼80 mins for each experiment on a standard computer with ten physical cores.

A total of 10,040 cell cycles from 784 lineages were used to statistically analyze their behaviors. Outliers within each cell cycle were ignored when calculating those time-dependent properties including sizes and elongation rates. Aberrant cell cycles such as filamentation were excluded from each experiment. Groupings of cell lineages according to their relative moments (values from 0 to 1) of CS onset within G0 lifespan were employed. Groupings are centered at intervals of 0.1, and each group encompasses a range of ± 0.05 around the center value.

### The allocation of protein synthesis during CS

The allocation of protein synthesis is extracted from the published protein abundance data^5^. In brief, a certain protein’s abundance equals its length multiplied by its number at sampling time. And total protein abundance is the sum of them. The allocation of protein synthesis to given proteins is calculated as the ratio of the change in their abundance to the change in total protein abundance between consecutive time points. Specifically, for 37 ℃ condition, protein synthesis allocation is directly equal to the corresponding protein abundance. This analysis focuses on the ribosome and CSPs (37 °C to 10 °C). Ribosome allocation is derived from the sum of allocations to its constituent ribosomal subunits. And the CSPs include the most highly induced 45 genes after CS^5^. Filtered genes in the original paper are not included. Although the duration of acclimation varies with temperature (∼2 h at 14 °C, ∼4 h at 12 °C, and ∼6 h at 10 °C), the corresponding increases in total protein synthesis are comparable (13% at 12 °C in this study; 12% at 10 °C from Ref. 15). To ensure consistent comparison across temperatures, we selected three standardized time points to illustrate: before CS (37 °C), mid-acclimation (3 h), and the end of acclimation (6 h).

### Experiments of batch culture

Both *E. coli* MC4100 and MG1655 strains were measured (MC4100 is presented in the text unless otherwise stated). A few single colonies from a Luria-Bertani agar plate were picked. Cells were cultured overnight at 37 ℃ in RDM and then placed in a thermostatic shaking water bath at 37 ℃ after subculture three times (> 10 mass doublings). The bacterial population was proven to reach a steady-state by then^63^. The measurements started when the cells were diluted again to an OD of 0.02 at 600 nm, with a sampling interval of 15-30 minutes. The OD_600_ value was determined using a spectrophotometer (Thermo Fisher Scientific, Genesys 150), with an average of three readings taken. 50 µl of the cell suspension was taken and fixed with the 450 µl of precooled fixation buffer (0.12% formaldehyde with 0.9% NaCl, filtered with a 0.22 µm filter), then used to measure the cell concentration and volume through a flow cytometer (Beckman Coulter, CytoFLEX S) in three parallel samples (Supplementary Note 5). Average cell volume was determined by the median of the forward scatter signals (FSC-A). A variant method called axial light loss detection is employed on the cytometer to detect FSC intensity, which is directly proportional to the cell volume regardless of the refractive index and thus benefits^64^. Further quantitative research also supports the relationship^63^. For the experiments with CS, when OD_600_ reached about 0.2, the culture temperature was gradually lowered to 14 ℃ or 12 ℃ within five minutes by mixing ice into the water bath.

### Total protein quantification

Total protein quantification was carried out according to previous studies using the Biuret method^65,66^. We quantified changes in total protein to represent Δ*M* since *M* is roughly proportional to the protein amount^67^. The MG1655 strain was selected and followed the culture procedure. The temperature was lowered to 12 ℃ to ensure a sufficiently long acclimation (∼5 hours) and higher resolution. Two sampling time points were at the beginning and near the end of the acclimation, respectively. Cell cultures were collected by centrifugation in triplicate (4 ml each) at each time point. The cell pellet was rinsed with 1 ml 0.9% NaCl solution, re-suspended in 0.2 ml water, and snap frozen in liquid nitrogen. The cell pellet was then thawed in a room temperature (RT) water bath. To extract proteins, 0.1 ml 3M NaOH was added, incubated at 100 ℃ heat block for 5 min, and cooled in water bath at RT for 5 min. Then 0.1 ml 1.6% CuSO_4_ was added to the sample, shaken and mixed thoroughly, and allowed to stand for 5 min. The sample was centrifuged and OD value at 555 nm was read by the spectrophotometer. The protein amount of the sample was determined from the standard curve of bovine serum albumin obtained on the day of the experiment using the same procedure.

### Simulation with the Rayleigh-Gans approximation

The intraparticle interference can be corrected by introducing a form function *P*(*θ*) to the Rayleigh differential scattering cross section *σ_d_*. If the relative refractive index of the scatterer in the surrounding medium (*m* = *n_s_* / *n_m_*) approaches to 1, *σ_d_* can be simplified as:

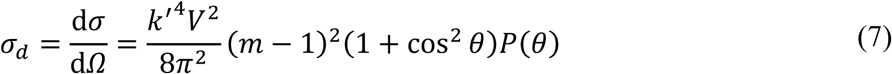

where *k’* is the wavenumber in the medium, *V* is scatterer’s volume, and *θ* is angle of observation. The refractive indices of *E. coli* and the surrounding medium are estimated as 1.384 and 1.338 at illumination wavelength of 600 nm^68^, respectively, thus meeting the prerequisites of the above equation. Furthermore, the refractive index difference between the scatterer and the medium is proportional to the dry-mass density *ρ_dry_*, with a scaling factor of the specific refraction index increment^68–70^, hence the quantity (*m* – 1) can be substituted by *ρ_dry_*. The *P(θ)* for differently shaped scatterers have different analytical forms. Ellipsoid of revolution as the form function is adopted to describe single *E. coli* cell, which has the following form:

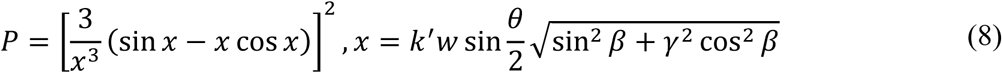

where *β* is the orientation angle between the ellipsoid and the direction bisecting the incident and the scattered light beam, *w* is width, and *γ* is aspect ratio. For randomly oriented bacteria in the medium, the form function should be averaged over all equally probable orientation angle *β*. Over a thousand cells with appropriate parameter ranges, were simulated to compute the total scattering cross section *σ* by integrating *σ_d_* over all angles except the detecting regime. Equivalent volume and aspect ratio were chosen as two independent determining parameters (ranging from 0.5 ∼ 5 µm^3^ and 2.5 ∼ 5, respectively) to simulate cells. To account for the uncertainty in the detection angle, we combined a series of integrals with lower bounds on *θ* varying from 0.5° to 3° at 0.5° interval and upper bounds of 180°.

## Data availability

All the data used in this study are available upon reasonable request to the corresponding author.

## Code availability

All codes are available upon reasonable request to the corresponding author for nonprofit academic research use only. Codes for controlling the LUNA-enabled bespoke platform and analyzing raw images are available in executable forms.

## Acknowledgements

We thank H. Cao for providing MG1655 strain. We thank H. Zheng for discussions. This work was supported by the National Key R&D Program of China (Grant No. 2024YFA0919600), Strategic Priority Research Program of the Chinese Academy of Sciences (Grant No. XDA0510200), National Natural Science Foundation of China (Grant No. T2525031).

## Author contributions

S.L. devised LUNA, developed the optical model and numerical simulation, performed the implementation and evaluation of LUNA, devised and implemented the LUNA-enabled bespoke platform, developed the standalone control software, developed image processing algorithm MARCEA, developed the scattering model and performed simulations on OD measurements, and analyzed all the experiments data. Z.M. participated in the evaluation of LUNA, conceived the biological experiments, prepared the bacteria samples, performed the biological experiments, and participated in the biological data analysis. Y.Y. designed the microfluidic chip, conceived the biological experiments, and participated in the biological data analysis. J.W. and Y.S. prepared the bacteria samples and performed the biological experiments. X.C. participated in the discussion of optical principle and the evaluation of LUNA. X.F. participated in the discussion of biological experiments. S.H. conceived the whole project and supervised the research. S.L. wrote the paper with contributions from all of the authors.

## Supplementary Information

is available for this paper.

## Correspondence and requests for materials

should be addressed to Shuqiang HUANG.

## Supplementary Information

### Supplementary Note 1. Current autofocusing methods

Focus drift can cause defocus and impede high-quality microscopy imaging. It comes from a variety of factors such as mechanical vibrations, thermal fluctuations, sample or stage movement, etc. An autofocus system detects and quantifies the defocus distance, which is then being send to the objective actuator to iteratively compensate until the optimal focal plane of interest is reached. Current autofocus techniques fall into two main categories: imaging-based methods, which capture a series of images at different planes along the z-axis and analyze the image quality to determine the focal position, reflection-based approaches, which employ auxiliary light to directly measure the distance changes between the objective and sample.

The imaging-based group usually employs varieties of image processing algorithms to characterize a focus score (such as edge sharpness, intensity contrast, entropy, etc.) of the captured image to evaluate the focus quality. The performance of those image algorithms based on diverse principles depends on the imaging modes and specimen, thus limiting their general applicability^1,2^. The optimal focal position is determined by finding the maximum of the focus score in z-stack images along the optical axis. However, the scanning process of image acquisition for z-stack is time-consuming, which may lead to photobleaching or phototoxicity to biological samples. Several ways to address this issue had been developed by lowering the requirement of accuracy: (a) Using fitting models to estimate the focus position, fewer images are required^3,4^; (b) An interpolated focus map created prior to the image acquisition is helpful in whole slide imaging, which is considered a disruptive technology in pathology practice^5,6^; (c) Direct measuring the defocus distance from images without z-scan, focus position can be determined by additional information such as phase detection (typically adopted in digital cameras) and deep learning^7–10^. A natural drawback among the image-based methods is that the focus score is insensitive to the focus change within the depth-of-focus (DOF) of the objective, thus limiting the accuracy of focus correction. Such methods would also fail to determine the focal plane when every image remains clear regardless of the axial position within a wide range, such as confocal microscopy. In addition, the focusing range is constrained due to the lack of image details as the defocus distance increases^11^.

The reflection-based group usually employs an external light source (typically a light-emitting diode or laser) to detect the axial position of a reflection interface. The light beam is reflected by this interface (usually the air-glass or glass-specimen with oil immersion objective) and collected by the optical detection system. As the reflected beam is sensitive to the axial disturbance of the focal plane, the defocus distance can be determined accordingly. Many implementations that utilize diverse signal detection schemes are demonstrated in laboratories. An intuitive and straightforward way to achieve accurate autofocusing is searching for the maximum reflected intensity of a size-limited light beam, yet axial scanning is required^12,13^. The defocused beam pattern can be used to locate the reflection interface without axial scanning. The detection optics can be modified by multiple detectors or astigmatic optics to measure the defocus distance when the beam is coaxial, whereas optical aberrations limit the detection sensitivity^14–16^. Besides, by introducing oblique illumination, which is also widely adopted by most commercially autofocusing products (For example, PFS from Nikon Instruments, AFC from Leica Microsystems, DF2 from Zeiss Microscopy, etc.), to generate asymmetric patterns, of which the centroid and intensity distribution are detected to identify the defocus status^17–19^. Precise alignment is required in those complex optical systems^20^, making a low tolerance of system performance to the mechanical errors in the optical train. Although the focusing range up to several times the DOF was achieved^14^, it is limited by the rapidly decreasing intensity of the defocused pattern. Another ubiquitous issue arises in the reflection-based methods, additional movement that reduces the focusing accuracy and precision after the repeated focusing procedure is necessary to compensate for an offset distance between the desired focal plane and the reflection interface^21^.

### Supplementary Note 2. Simplification of coma-induced optical model

In discussing the light distribution near the focus of an optical system, we shall consider a Gaussian beam as the monochromatic light source. In this note, we follow the notations in main text. The incident beam location is set to relative polar coordinates (*ρ_0_* = 0.1, *θ_0_* = 0°) in the objective pupil plane (Note 2. Figure 1). The coordinate system is translated to the beam center as (*r*, *φ*).

**Note 2. Figure 1.**
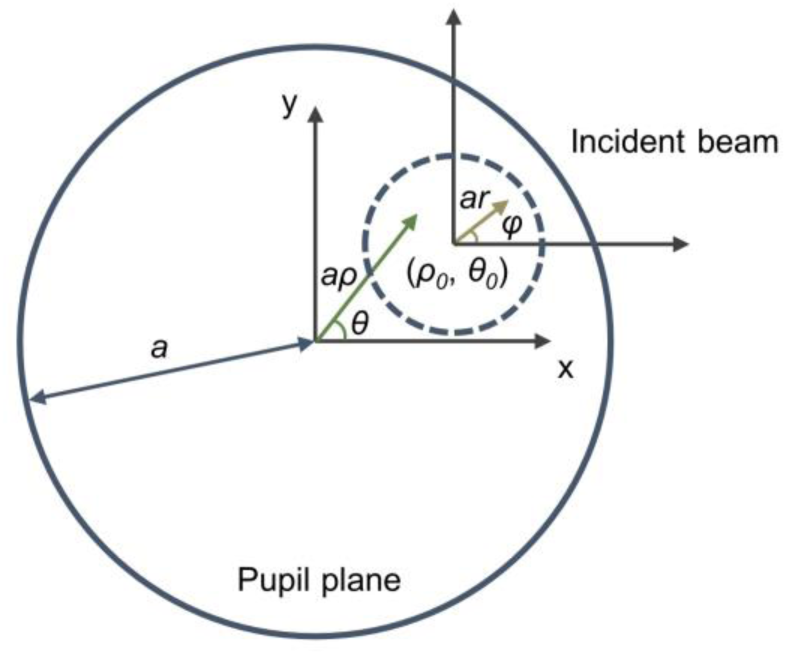
Incident beam at the pupil plane. The radius of the objective pupil is *a*. The dashed circle indicates the incident beam. The origin of polar coordinates (*ρ*, *θ*) is the pupil center. The origin of polar coordinates (*r*, *φ*) is the incident beam center (*ρ_0_*, *θ_0_*).

Applying the Huygens-Fresnel principle, the complex amplitude of point *P* is obtained by:

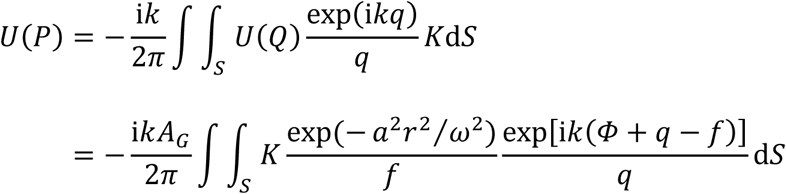

The inclination factor *K* is equal to 1 by assuming the angle between the normal of the primary and the secondary wavefront is small. Accordingly, the off-axis amount of the incident beam is considered to be a small value, therefore the variation of *K* over the integral area can be omitted. In the bespoke microscope system, the distance *OP* is small compared to *f*. Therefore, *q* is replaced by *f* in the denominator of the integrand, and *k*(*q* – *f*) is approximated in the Cartesian axes at *O*:

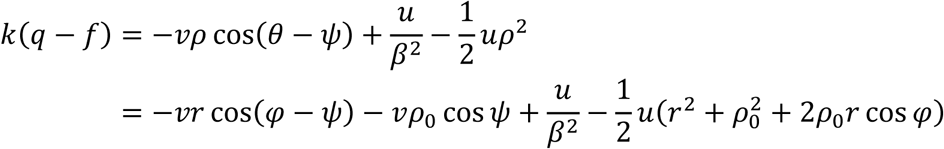

Combining the phase terms lead to:

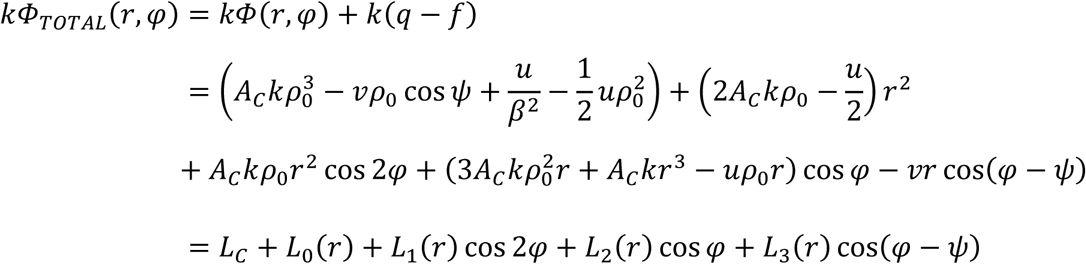

Herein, we consider the dominating lower order coma (or primary coma) aberration. The coma aberration function can easily be expressed by *Φ*(*ρ*, *θ*) = *A_C_ρ*^3^ cos*θ*, where the coefficient *A_C_* is proportional to the field angle (angle between the chief ray and optical axis). Further, the element is d*S* = *a*^2^*r*d*r*d*φ*. Hence, *U*(*P*) becomes:

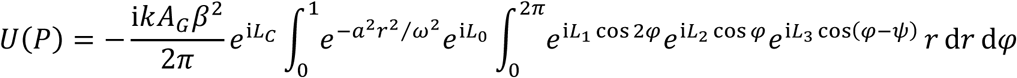

The above integral can be expanded into an infinite series with the help of the Jacobi-Anger identity:

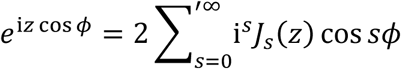

Note the term *s* = 0 is taken with a factor 1/2, and we adopt this configuration in the following summation sign with a prime. Multiplying the expansions of the integrand with angle *φ* we get:

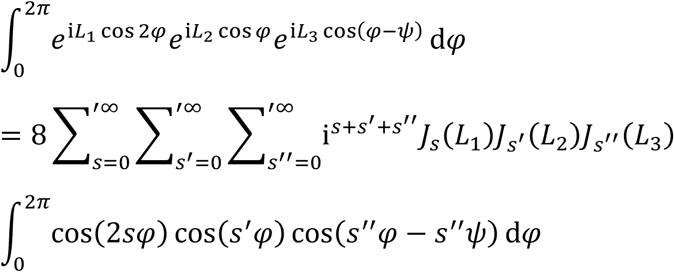

The integrand with the form of [cos(*Aφ*) cos(*Bφ*) cos(*Cφ-Cψ*)] is not zero only in limited cases. In addition, we introduce the Graf’s addition theorem for Bessel functions before the simplification:

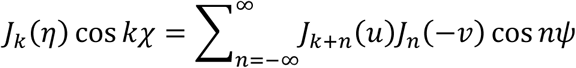

where *η* = (*L*_2_^2^ + *L*_3_ ^2^ + 2*L_2_L_3_* cos*ψ*)^1/2^, tan*χ* = -*L_3_* sin*ψ* / (*L_2_* + *L_3_* cos*ψ*). Hence, expanding the summation and ignore the zero terms, the integral is simplified as:

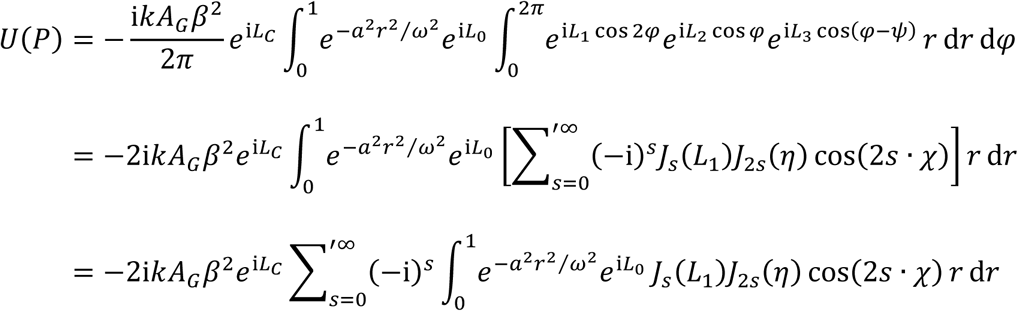

In this way, the original double integral is reduced to the summation of an infinite series of integrals. In fact, it is only necessary to take the first four terms of the summation series to obtain adequate intensity accuracy (Note 2. Figure 2). For those scenarios where precise intensity is not required, three terms will suffice. In this work, we select the first four terms to effectively calculate the intensities.

**Note 2. Figure 2.**
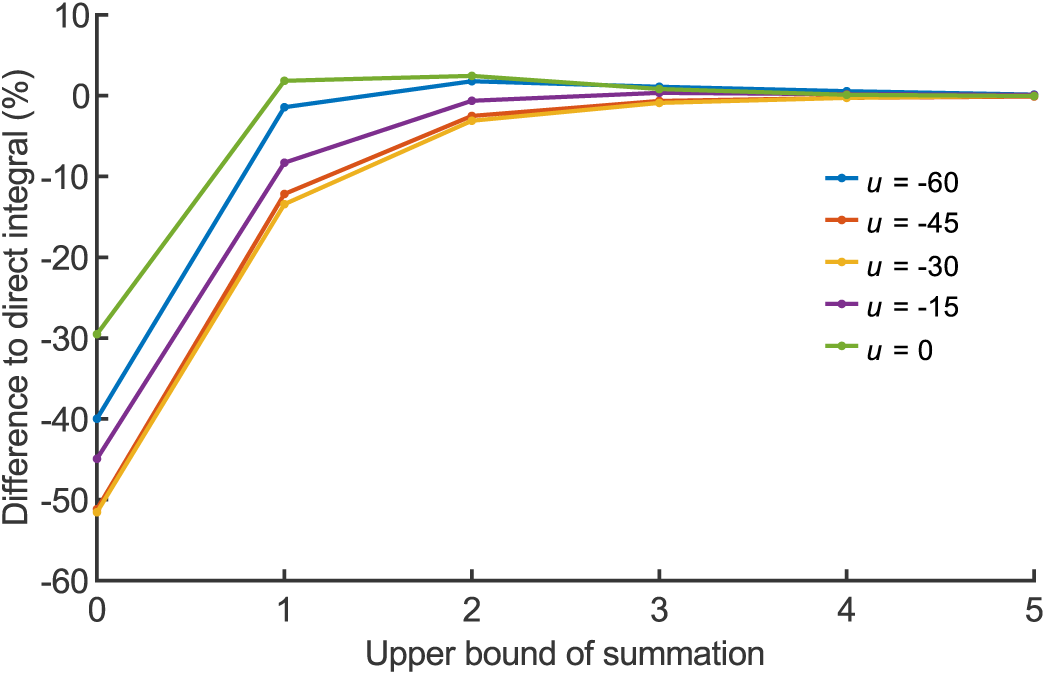
Convergence of intensity calculation with increasing series terms. Relative differences (%) compared to direct calculation of the double integral is shown. Five locations (*u*) on the optical axis were chosen.

Furthermore, this simplification saves a considerable amount of time compared to the time taken for the double integral. We used MATLAB to evaluate the time cost. Parallel computing with four physical cores was enabled for both evaluation conditions (on a workstation with CPU @ 3.70 GHz). We tested intensity distribution on the focal plane with different pixel sizes. The average time cost with a standard deviation of three repetitions is shown (Note 2. Figure 3). As the computational scale increases, the time consumed by directly computing the double integral is tens to hundreds of times that of the simplified method. For reference, computing an intensity distribution of 800×1000 pixels (e.g. the x-z plane in main text Figure. 1d) takes ∼718 seconds using the simplified method.

**Note 2. Figure 3.**
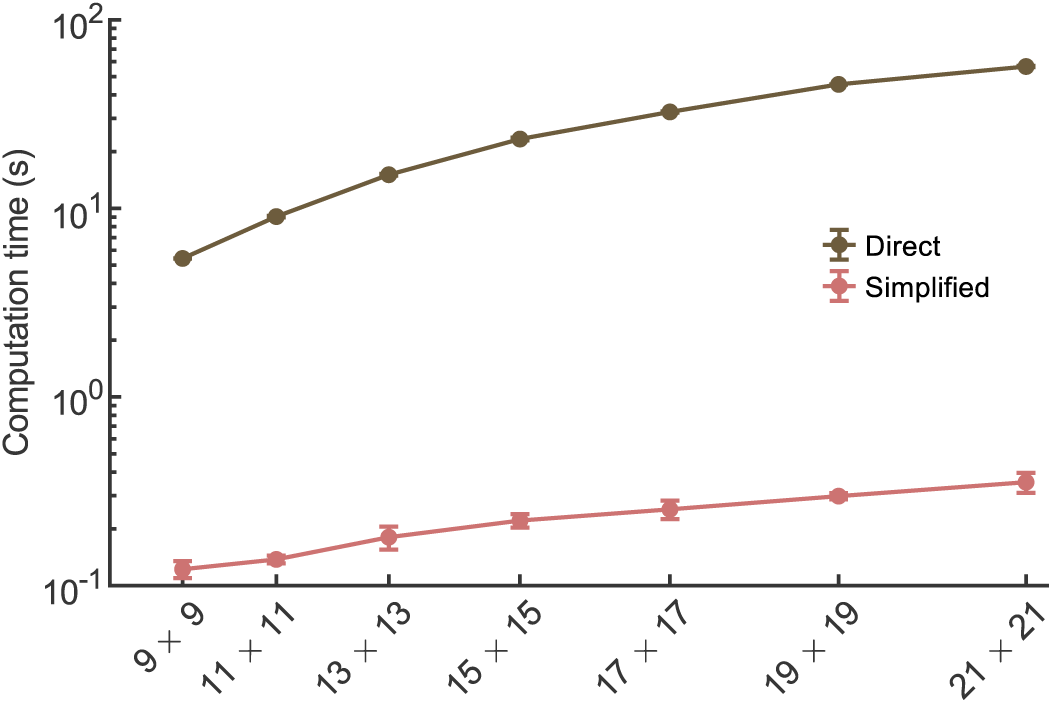
Computation time comparison between direct and simplified methods for varying pixel size. Lines, average time cost; error bars, standard deviation.

Apparently, the coefficient *A_C_* in the primary coma aberration function is determined by the oblique angle of the incident light beam. The variation range of *A_C_* will not be significant, as it is limited by the lens parameters used in the optical model. Larger *A_C_* results in a more extended range with a high contrast pattern, allowing LUNA a longer working range (Note 2. Figure 4, see also main text Figure. 1d). However, this also reduces the change rate of the brightest spot’ centroid along the optical axis, which reduces LUNA’s sensitivity.

**Note 2. Figure 4.**
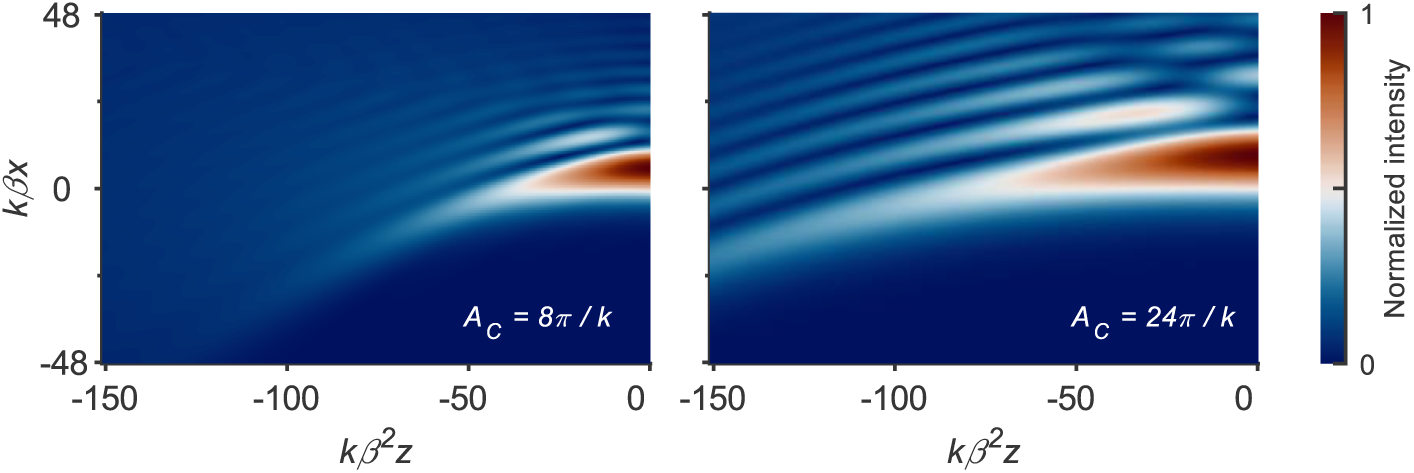
Computed intensity distributions corresponding to different *A_C_* values. Image contrast was adjusted for better exhibition (Gamma = 0.5). *k* is wavenumber, *β* = *a* / *f*.

### Supplementary Note 3. Single cell image analysis

#### Part 1: Cell identification algorithm pipeline

Time-lapse imaging of cells generates large amounts of data. In order to obtain detailed morphological information of individual cells, fast and reliable automated analysis is inevitable. We developed an automated image processing software whose core algorithm is named MARCEA (Microfluidic Architecture based Resilient Contour Expansion Analysis) to address this problem (Figure S11a). The raw input for MARCEA can be either fluorescence or phase-contrast images.

In this work, MARCEA targets bacterial growing in a microfluidic chip known as mother machine. This chip contains multiple microchannels, with one open end for the growth of the trapped cells. The growth medium diffuses from the connected main trench (about 25 µm in height and 100 µm width) into the microchannels. The field of view (FOV) of each imaging site covers approximately 25 microchannels. The raw image stack consists of time-sequential images of multiple imaging sites. To extract individual microchannels in the raw phase-contrast image, a preprocessing procedure including serial tasks is required, as follows:

a. Correct orientation. The raw image is first binarized to segment the microchannel objects from background. The rotation angle and channel direction are then determined by fitting the edges of the microchannel closed ends. The image is rotated and flipped to orient the microchannels vertically with the open end on top. Bilinear interpolation method is adopted since the rotation angle is usually small.
b. Select region of interest (ROI). Cells near the exit of the microchannels are pushed out as cells grow and divide within the microchannels and therefore ignored. The rotated images are cropped from the bottom to near the top of the microchannels to avoid redundant data.
c. Locate channel stripe. A duplicate of the ROI image is used to locate the vertical edges of each microchannel. This duplicate is then denoised and the background equalized. The precise coordinates of the edges are determined by the horizontal intensity profile of the duplicate. The stripe images of the microchannels can be extracted individually from the ROI image thereafter.
d. Segment channel region. The closed end of the microchannel is designed to be semicircular. Unlike detecting the vertical edges, cells are most likely attached to the round ends, resulting in smaller intensity differences. Therefore, the gradient of the stripe image is computed using the Sobel operator to estimate the rounded shape. Smoothing the maximum gradient boundary is followed to determine the fine edge. The entire microchannel region is constrained by the vertical and bottom boundaries.

The ROI image is further processed to identify cell contours. An improved Otsu method is applied to each image stripe of the microchannel to estimate a rough sketch of the cell objects. In the experiments of this work, we focus only on the cells at the bottom of the channels (mother cells). A skeletonization process is applied to extract an initial guess for the centerline of a binarized cell object. Starting from the initial centerline, resilient expansion is iteratively executed to fit the object’s shape to a cell model. This model for *E. coli* cells is defined as a slightly curved rod shape. After each iteration step, the centerline is modified to adapt the updated binarized object. The iterations stop when the signal-to-noise ratio of the detected boundary reduces below a predefined threshold. The above process will give us candidate mother cell, waiting for split detection.

The split detection is based on the detection of width and peak intensity along the centerline. Some criteria are applied to filter false split sites caused by noise fluctuations. If any split site is detected from the candidate cell, segmentation is performed and the precise boundary detection process is repeated for the split cells. Once the mother cell is identified, its accurate morphological features are recorded for statistical analysis (e.g., area, centroid location, centerline coordinates, width along centerline, etc.). The same process could also be applied to all the cell candidates in the microchannel if needed.

In the implementation of MARCEA, some basic morphological functions were optimized, greatly improving the image processing speed. The computational performance was evaluated in 10 parallel processes in two scenarios:

|  | Workstation 1 | Workstation 2 |
| --- | --- | --- |
| CPU (Intel) | Core (TM) i9-10900 @ 2.80 GHz | Xeon (R) W-2255 @ 3.70 GHz |
| Physical cores | 10 | 10 |
| RAM | 32 GB | 64 GB |
| Data location | External hard drive disk | Internal hard drive disk |
| Dataset | 44,910 frames from 30 positions and 1497 time points, ~115 GB | 30,780 frames from 27 positions and 1140 time points, ~90 GB |
| Total processing time | 48 min 32 s | 27 min 51 s |
| Speed (frames per second) | ~15.4 | ~18.4 |

#### Part 2: Cell tracking automation

The main target of cell tracking is to link the same cells in consecutive frames. The prerequisite step is to calibrate the lateral frame shift, which is detected via cross-correlation. This is typically robust to small shifts during normal time-lapse imaging. The vertical orientated microchannels can be easily aligned. However, under cold shock, the lateral shift between consecutive frames may reach ∼10 µm (Figure S11b), across several microchannels. And this can lead to microchannel mismatch. To deal with this situation, cell features (including centroid coordinates, smallest box containing the cell region, length, orientation in the microchannel, etc.) are employed as unique characteristics of each microchannel. The maximum probability of the mismatch offset is determined by finding the minimum distance in the characteristic space.

A general assignment task for the mother machine chip usually involves solving a global optimization problem. Possible events for a cell include: unchanged (growing or moving), divided, lost (due to cell lysis or being flushed out of the microchannel), appeared (e.g., misidentification or, rarely, entry into the microchannel). Calculation of probabilities of each case takes into account the morphological parameters of the cell. The overall probability gives the division event time point. Manually inspection and correction for the division events was carried out for ground truth data.

With only the mother cells in each microchannel being studied, we ignore the other offsprings away from the bottom of the microchannels. The structural data matrix of mother cells is recorded for statistical analysis. The total time cost on the tracking is typically a few minutes.

### Supplementary Note 4. Theoretical OD dependency and numerical simulation

Rayleigh scattering applies to scatterers which are small compared to the wavelength of incident light. The differential scattering cross section is:

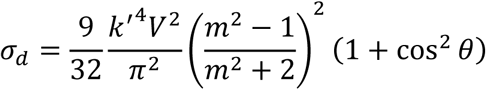

This equation applies when (*m* – 1) is small. For larger scatterers such as bacteria, more complicated treatment is needed. We use Rayleigh-Gans approximation by introducing a form function *P*(*θ*) to correct for the interference of the wavelets scattered from different parts of the bacteria. Most bacteria with high water content have low refractive index. We estimate the refractive indices of *E. coli* and the RDM (EZ rich defined medium) as *n_s_* = 1.384 and *n_m_* = 1.338, leading to *m* = 1.034. Therefore, *σ_d_* could be simplified to equation (7) in main text Methods. The total scattering cross section *σ* is obtained by integrating *σ_d_* over the entire scattering regime:

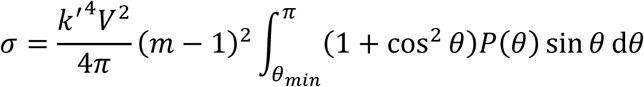

An ideal OD measurement would collect none of the scattered light. However, scattered light over the cone of a few degrees could still fall on the spectrophotometer’s detector (Figure S15a, S15b). In the expression of *P*(*θ*) exists an orientation angle *β* indicating the ellipsoid of revolution to the direction of the incident beam. To compute for randomly oriented cells, *P*(*θ*) is averaged over all equally probable orientations from 0° to 90°. A relationship of *σ* proportional to *V*^1.34^ is obtained by simulating *E. coli* cells with typical morphology parameters at an incident wavelength of 600 nm.

Furthermore, *m* could be related to the dry-mass density (*ρ_dry_*) of a cell, based on the assumption that *n_s_* is proportional to the cell’s dry matters:

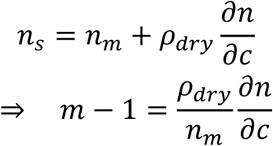

where ∂*n* / ∂*c* is the specific refraction index increment. Generally speaking, differences in composition in cells will produce little percentage differences in ∂*n* / ∂*c*, thus ∂*n* / ∂*c* can be assumed as constant^22^. Consequently,

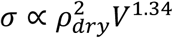

In turbidity measurement with incident light *I_0_*, OD is related to the transmission of light *I_T_*:

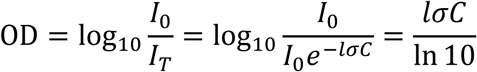

where *l* = 1 cm is the optical path length of the cuvette in the spectrophotometer. In the simulation performed, the average *σ* is 0.713 µm^2^, which leads to a scaling factor of 3.1E-9 ml from *C* to OD_600_. On the other hand, the same scaling factor in the bulk experiments without changing temperature (Figure S17b) is fitted as 2.6E-9 ml, which is close to our numerical simulation. Combining the constant coefficients into *α*, equation (2) in main text can be obtained at the given condition. The relationship holds at low concentrations of bacteria. In dense cultures, multiple scattering events may redirect the first cell’s scattered light into the detector, and the destructive interferences between cells must be considered^23,24^. This causes deviations from the Beer-Lambert law. The change in cell density and size also disrupts the relationship^25^.

The OD equation is also related to total dry mas per ml (*M* = *ρ_dry_VC* = *M_dry_C*) and single cell surface-to-mass ratio *R_SM_* = *S* / *M_dry_*:

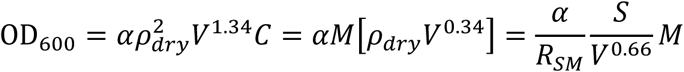

For *E. coli* cells under a steady-state, *R_SM_* usually remains nearly constant^26^. In the above simulation, bacteria were modeled as ellipsoids, and explicit *S* / *V*^0.66^ was calculated in the wide parameter space (Note 4. Figure 1). The coefficient of variation (CV) is 5.6%, indicating a stable constant value of the ratio *S* / *V*^0.66^. It’s worth noting that the parameter range of the simulated cells is quite extensive to cover different types of cells^27^, so the CV of this ratio is expected to be much lower for a single type of cell. In short, OD_600_ can only be considered as a total mass indicator at steady-state.

**Note 4. Figure 1.**
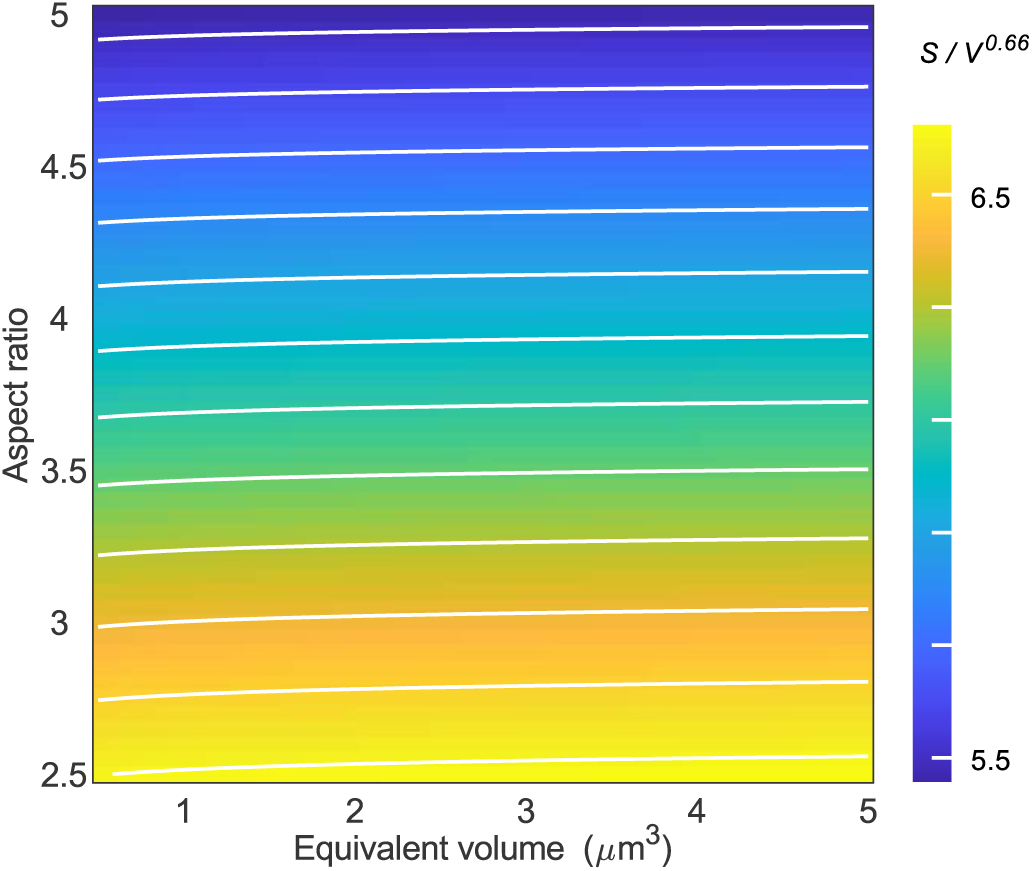
Calculated values of scaling ratio in the simulation. Color bar represents scaling ratio *S* / *V*^0.66^ for each cell. White contour lines indicate the elevation.

### Supplementary Note 5. Gating strategy used in flow cytometric analysis

Samples for cell counting, forward scatter signals (FSC) and side scatter (SSC) signals characterization were firstly fixed with pre-cooled cell counting buffer (0.12% formaldehyde with 0.9% NaCl). The fixed samples were diluted as necessary with staining buffer (0.9% NaCl with 1 µg/ml DAPI) prior to the flow cytometric analysis. Total cell number count in each measurement was controlled to range from 50,000 to 200,000. Gating strategies are illustrated below (Note 5. Figure 1).

In FSC-A/SSC-H plots, sample particles in P1 in panel **a** subtracted with control particles in P1 in panel **d** were regarded as the bacterial cells. Traditionally, DAPI staining is used to determine cells from particles. We checked that our gating strategy has no differences with it. For the selected particle subpopulation P1, plotting in DAPI-H/SSC-H (panel **b**) shows that it is equivalent to the particle subpopulation P2 selected directly under DAPI-H/SSC-H plots (panel **c**, **f**).

Samples at different time points have similar unimodal distributions in FSC-A/SSC-H plots (panel **a**) and bimodal distributions in DAPI-H/SSC-H plots (panel **c**). Considering the comparison between the two gating strategies (P1 and P2) for subpopulation classification, the unimodal distribution observed in the FSC-A/SSC-H plot allows for a more precise and consistent boundary definition. In contrast, the DAPI-H/SSC-H plot displays a bimodal distribution, which is more prone to variation over time. This variation could lead to challenges in defining the boundaries due to subjective uncertainties. Thus, the former gating strategy we adopted provides a more robust approach to reliably distinguishing subpopulations.

The gains for FSC, SSC and DAPI channels were set to 500, 500 and 2,000, respectively. The total particle counts of the subpopulations (P1 or P2) in panel **a**, **c**, **d** and **f** are 96148, 96275, 961 and 961, respectively.

**Note 5. Figure 1.**
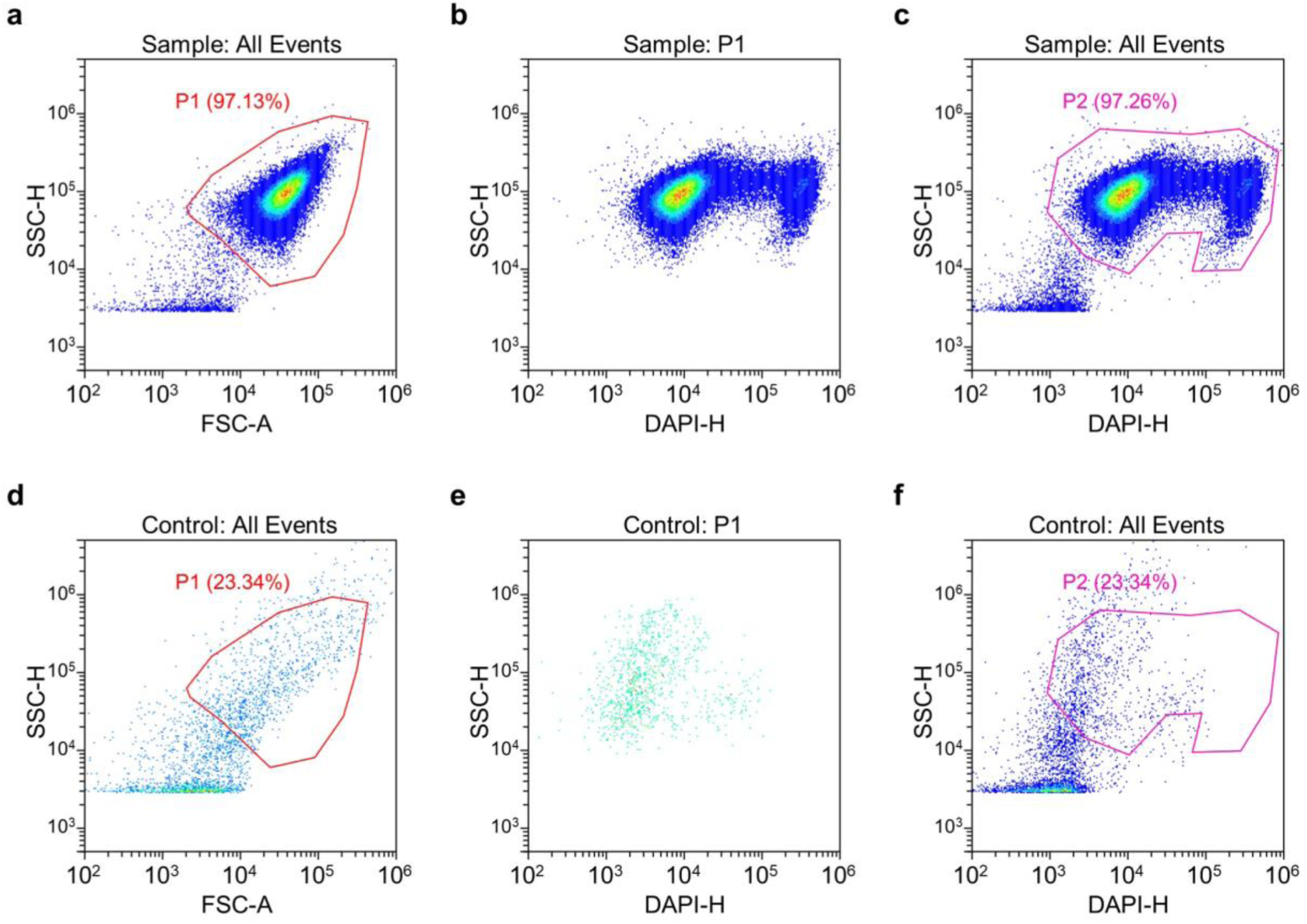
Gating strategy for isolating target cell population. Enclosed areas with colored lines: selected particle subpopulations, marked with data percentage.

---

**Figure S1.**
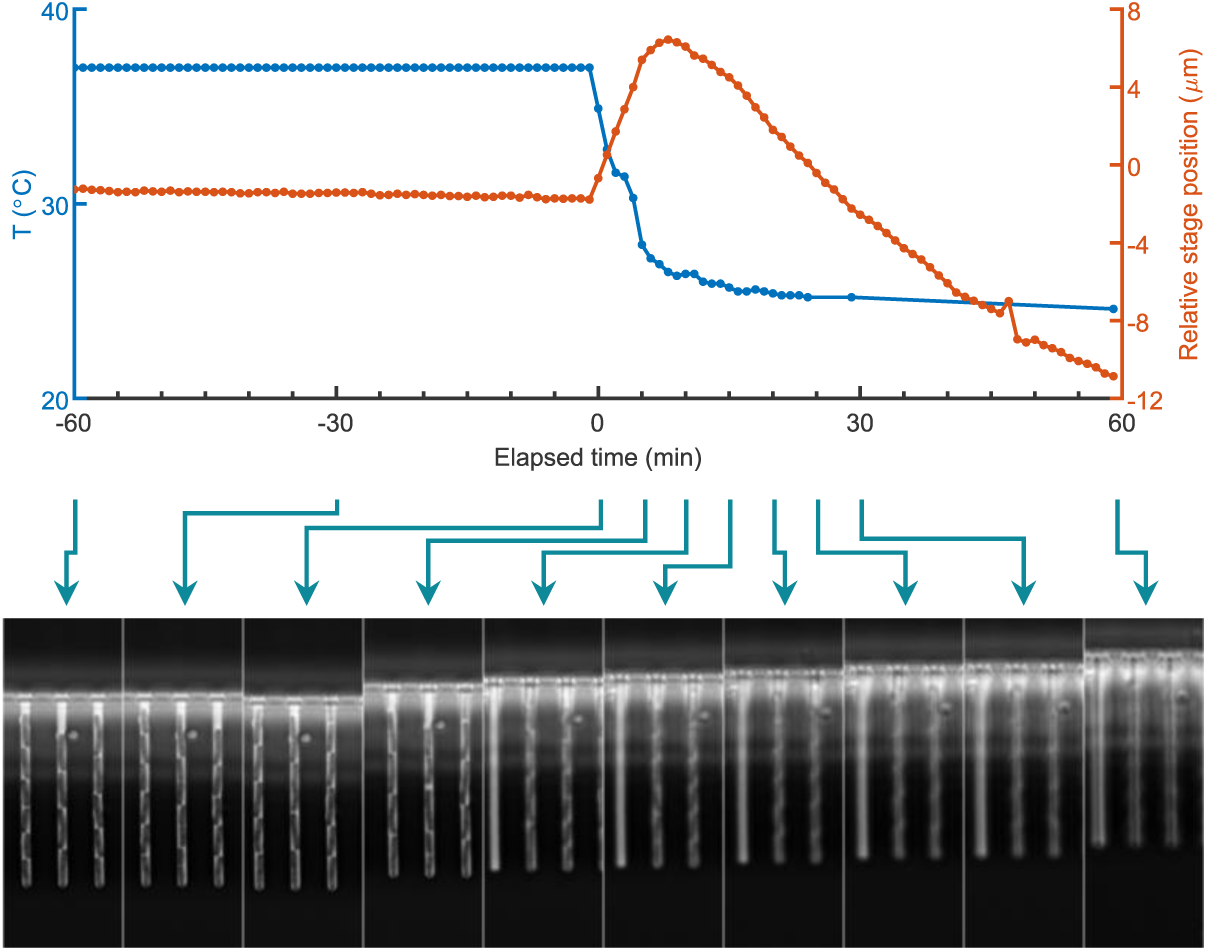
Time-lapse experiment with induced temperature downshift using commercial microscope. Time-lapse phase-contrast imaging of *E. coli* cells in a microfluidic chip (1 min interval, 20 position of views) was performed using a commercial microscope (Nikon Instruments, Ti2 with autofocusing unit PFS). The microscope stage and objective (100×, with lens heater) were enclosed in an incubator-style chamber with an external air-heating unit. Temperature was maintained at 37 ℃ until the start of cooling (*t* = 0), when the heating unit (and the lens heater) was turned off. The flow medium for cells was replaced with 10 ℃ medium, and the chamber cooled passively to room temperature (temperature stabilized at 25 ℃ after ∼60 min). The commercial autofocusing unit remain active throughout the experiment. Representative images from selected time point are shown.

**Figure S2.**
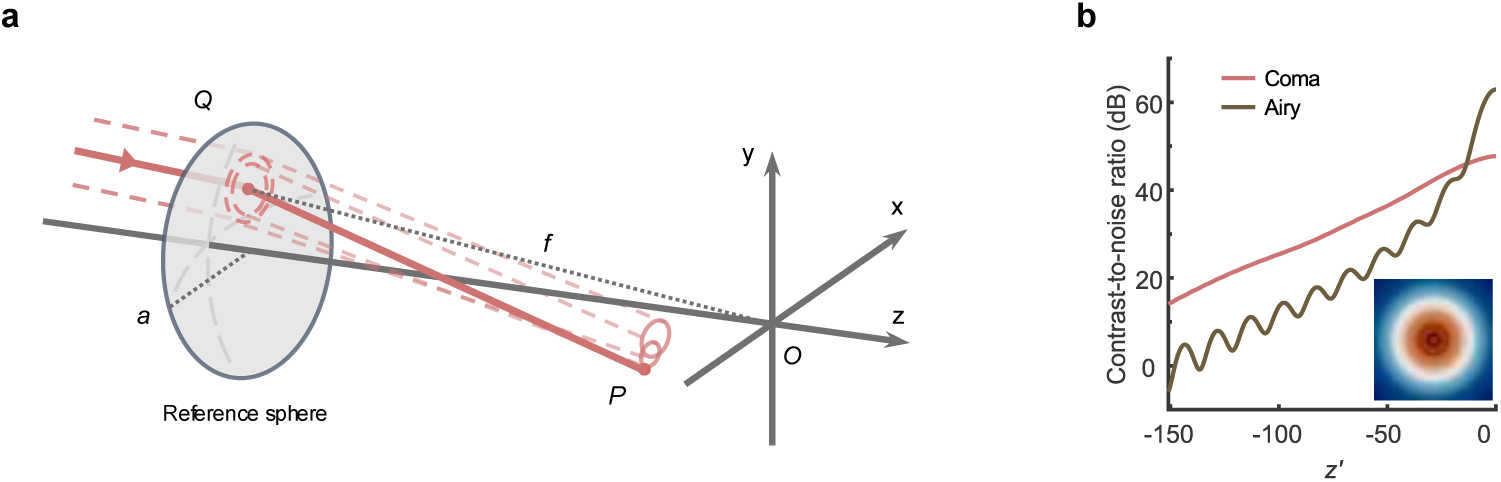
Optical model schematic and computed contrast comparison. **a**, Schematic diagram of the optical model. An off-axis Gaussian beam (red lines) propagates through an objective lens of radius *a* and focal length *f*. *Q*, point in which a light ray in the image space intersects the reference sphere that approximately fills the objective pupil; *P*, arbitrary point near focus. **b**, Comparison of spot contrast with and without coma aberration. The contrast-to-noise ratio metric, which influences the detection accuracy of the focus state, was introduced to evaluate the spot quality. Metric values were calculated from the computed intensity distribution images at different axial position. Coordinate *z’* = *kβ^2^z*, where *k* is wavenumber, *β* = *a* / *f*. Inset image: Airy pattern at -90 axial position in the dimensionless space in an aberration-free system.

**Figure S3.**
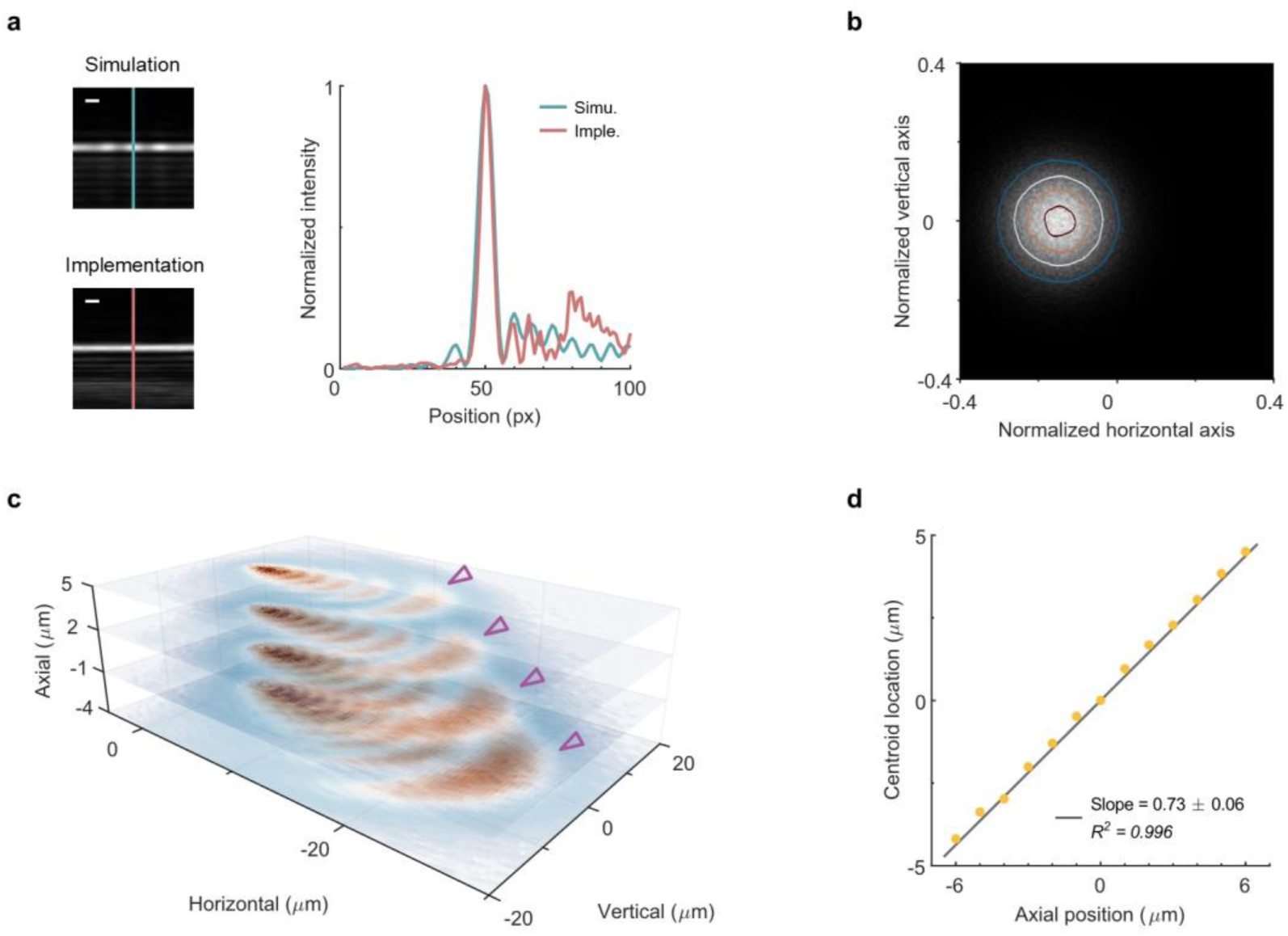
Ray tracing simulation of LUNA’s optics design. **a**, Comparison of numerical simulation and implementation. Left panel: Retrieved images from the detection planes in the simulation and LUNA implementation, respectively. Right panel: intensity profiles along the lines in both images. px, pixel. **b**, Intensity on the entrance pupil plane in the simulation. Polar coordinates were normalized by the aperture of the objective (100×). Colored lines indicate isophotes. Beam center distance is approximately *ρ_0_* = 0.15. **c**, Intensities on the reflection planes in the simulation. Images at each axial position were individually normalized. Triangle markers indicate the outermost crescent spots. **d**, Magnified defocus distance on the reflection plane. Linear regression with slope (95% confidence intervals) and fit goodness is shown.

**Figure S4.**
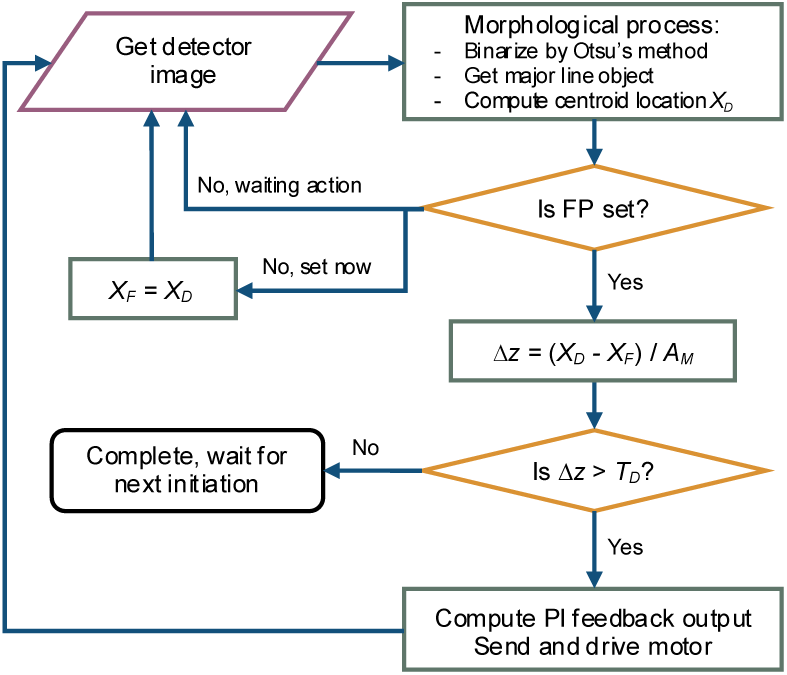
Flowchart of LUNA. FP, focal plane; *A_M_*, effective magnification from axial position to centroid location; *X_F_*, focal plane location; *Δz*, detected drift; *T_D_*, set threshold; PI, proportional-integral controller. Morphological process costs ∼2 ms for a single captured image from LUNA’s detector on the control computer with 3.60 GHz CPU.

**Figure S5.**
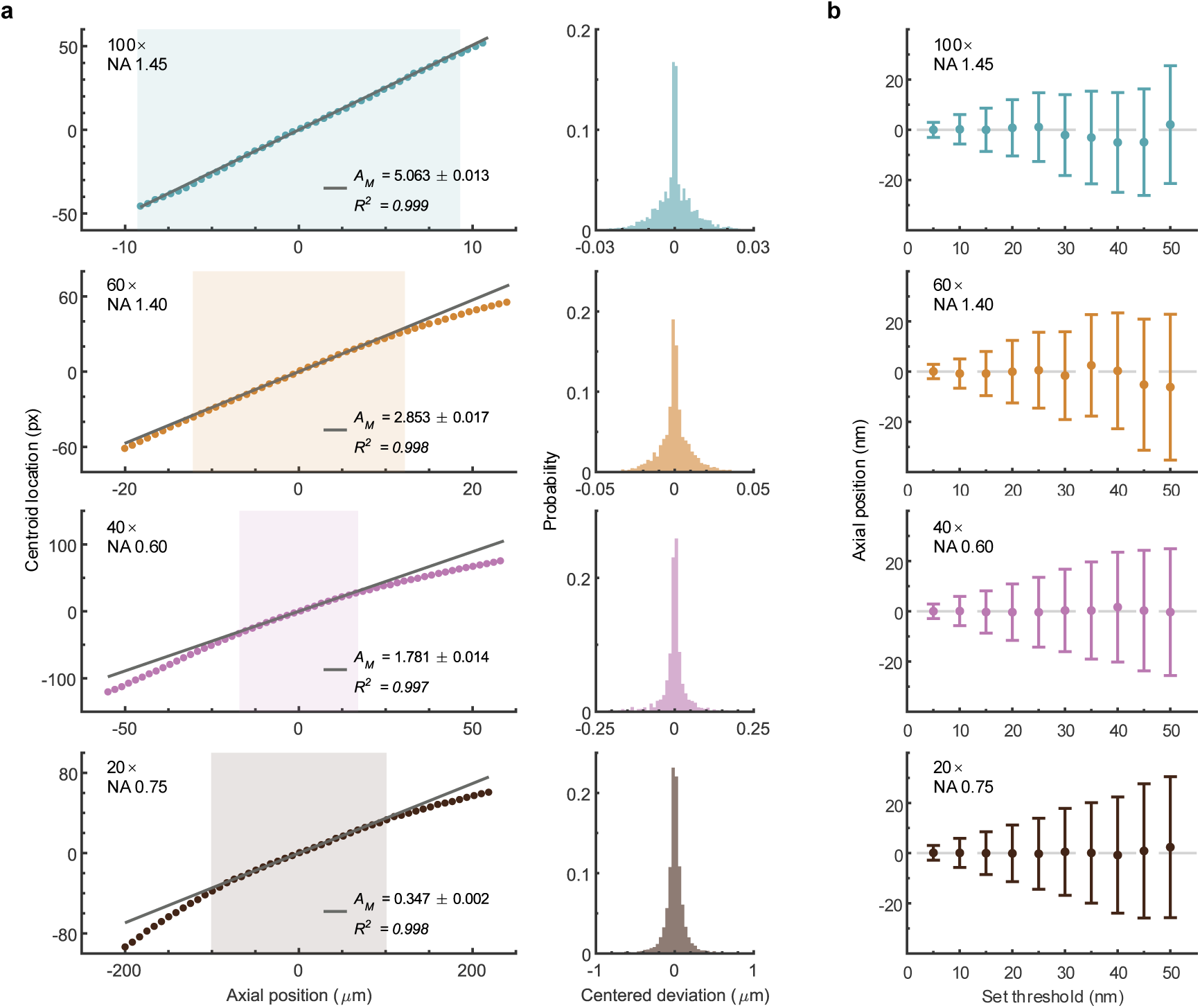
Supplemental performance metrics of LUNA. **a**, Calibration curve (left) and measurement error distribution (right) for representative objectives. Shaded regions in the left panels indicate the linear range of the corresponding objective. Linear regressions with slope (95% confidence intervals) and fit goodness are shown. **b**, Focusing accuracy and precision assessment. Dots and error bars represent the mean and s.d. values of the focus errors.

**Figure S6.**
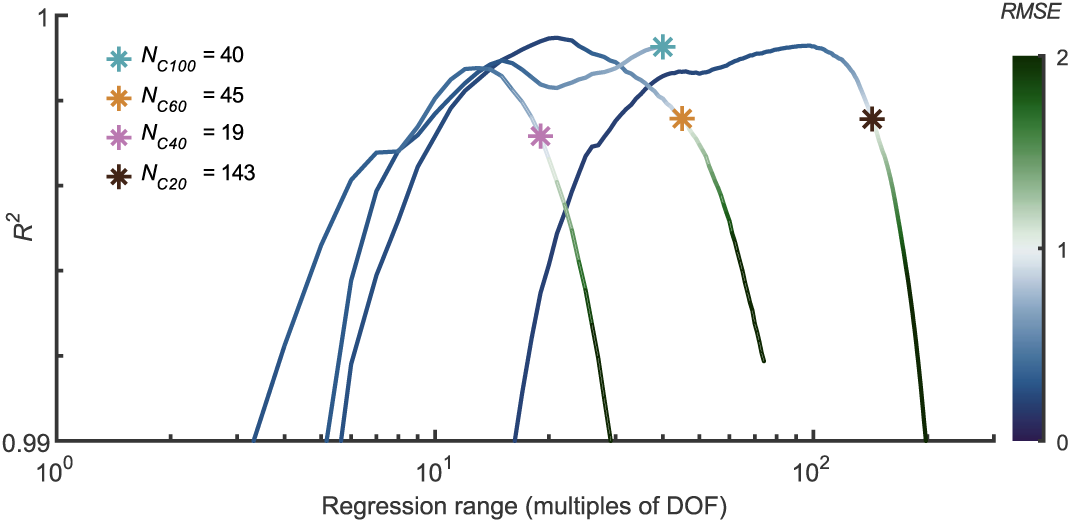
Evaluation metrics of linear regressions under different range. The regression range was symmetrically selected with the zero point as the center and multiples of the corresponding DOF. Critical *N* multiples of DOF was determined by restricting the fitting root mean squared error (*RMSE*) less than 1. Numbers in the subscripts indicates the objective magnification.

**Figure S7.**
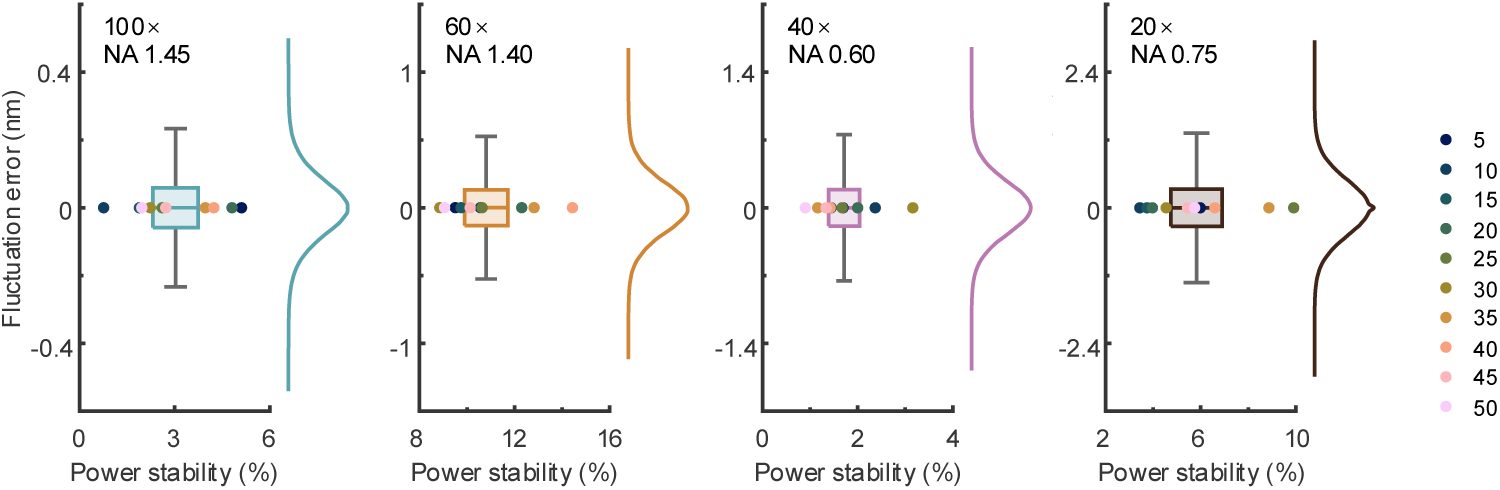
Impacts of intensity fluctuations on focusing accuracy. All data for the 10 set thresholds are presented as box plot with the edges at the 0.25 and 0.75 quantiles, the central line at the median, the width at the standard deviation, and the whiskers at the minima and maxima within 1.5× interquartile range from the box. Colored dots indicate average power stability of the 10 thresholds (right legend). Colored line, probability distribution.

**Figure S8.**
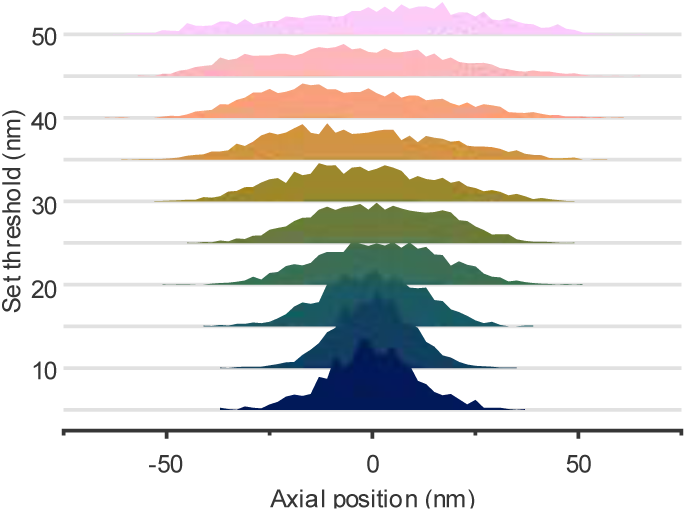
Ridgeline plot of relative drift after the previous autofocusing procedure. Drift distributions under different set thresholds at 1 Hz measurement. Colors indicate different set thresholds, unit in nm.

**Figure S9.**
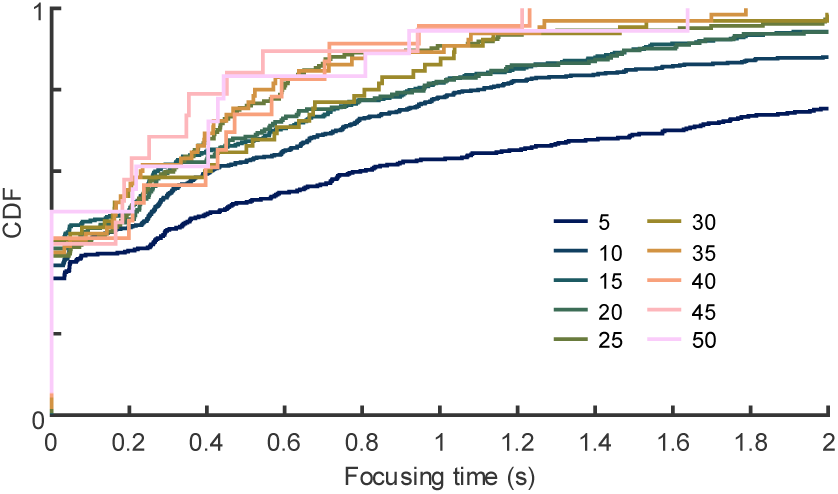
Cumulative distribution function of focusing time at 1 Hz measurement. CDF, Cumulative distribution function. Colored lines indicate different set thresholds, unit in nm.

**Figure S10.**
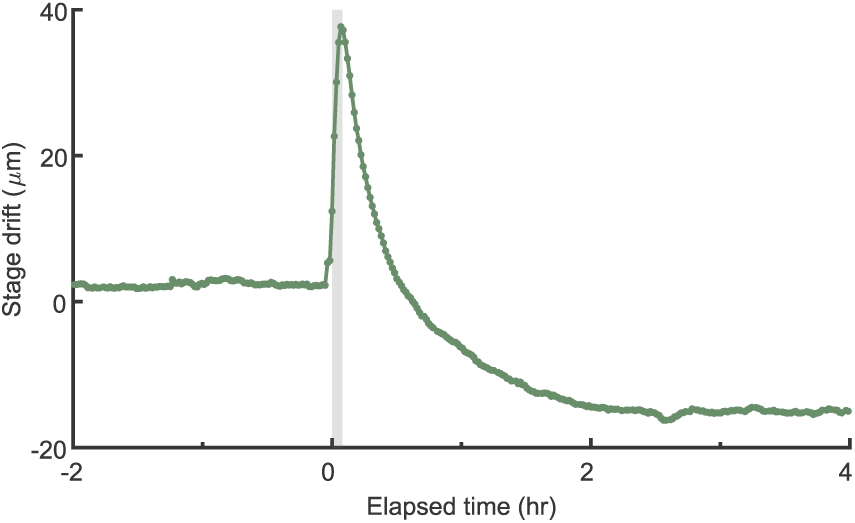
Stage drift over time. Data in scatter form represent the average of all imaged sites (*n* = 45). Shaded region indicates the rapid temperature drop period. Error bands, s.e.m.

**Figure S11.**
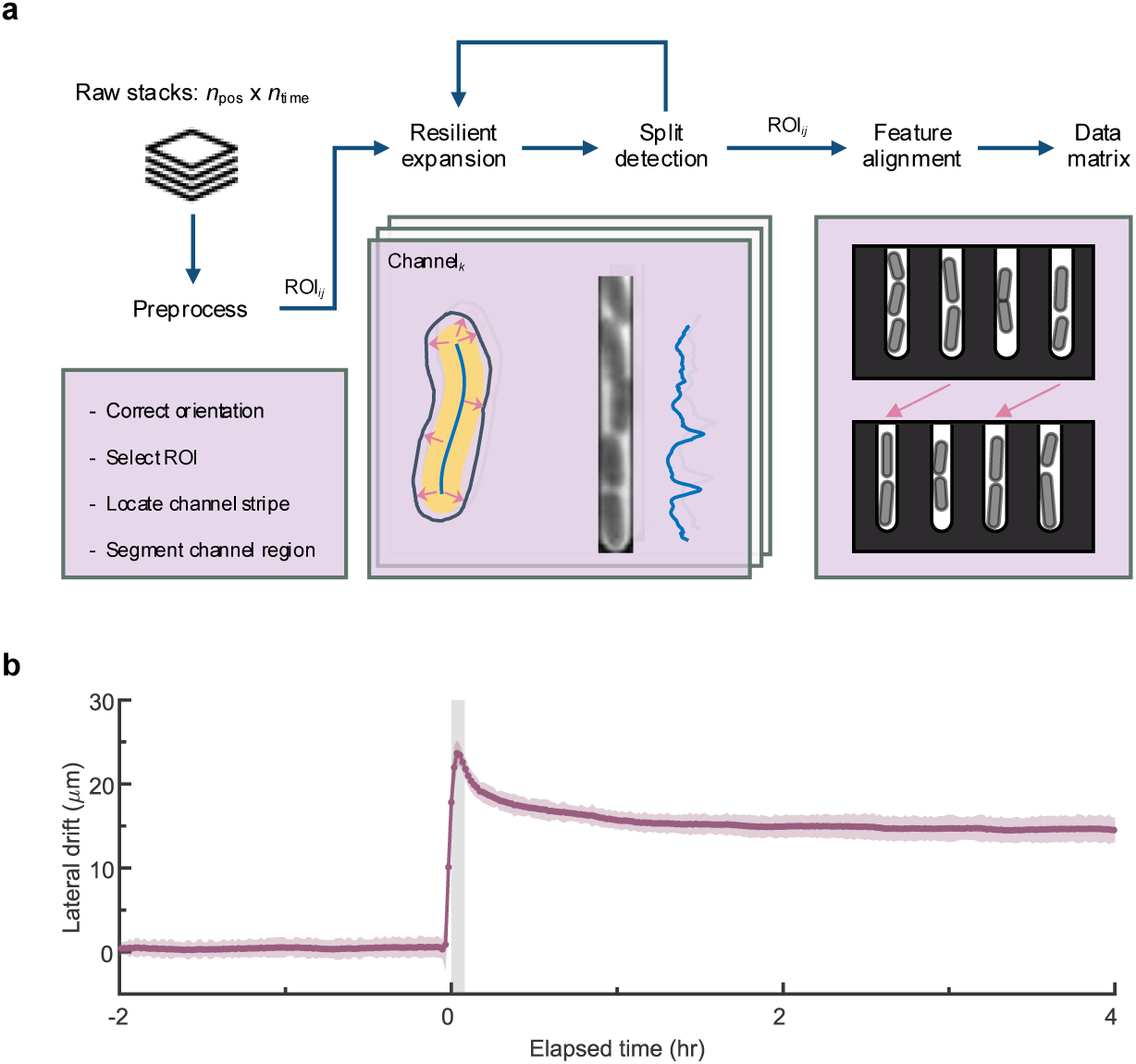
Quantitative image analysis and drift calibration. **a**, Microfluidic Architecture based Resilient Contour Expansion Analysis (MARCEA) pipeline. ROI, region of interest. **b**, Lateral drift of imaging sites over time. Data in scatter form represent the average of all imaged sites (*n* = 45). Shaded region indicates the rapid temperature drop period. Error bands, s.e.m.

**Figure S12.**
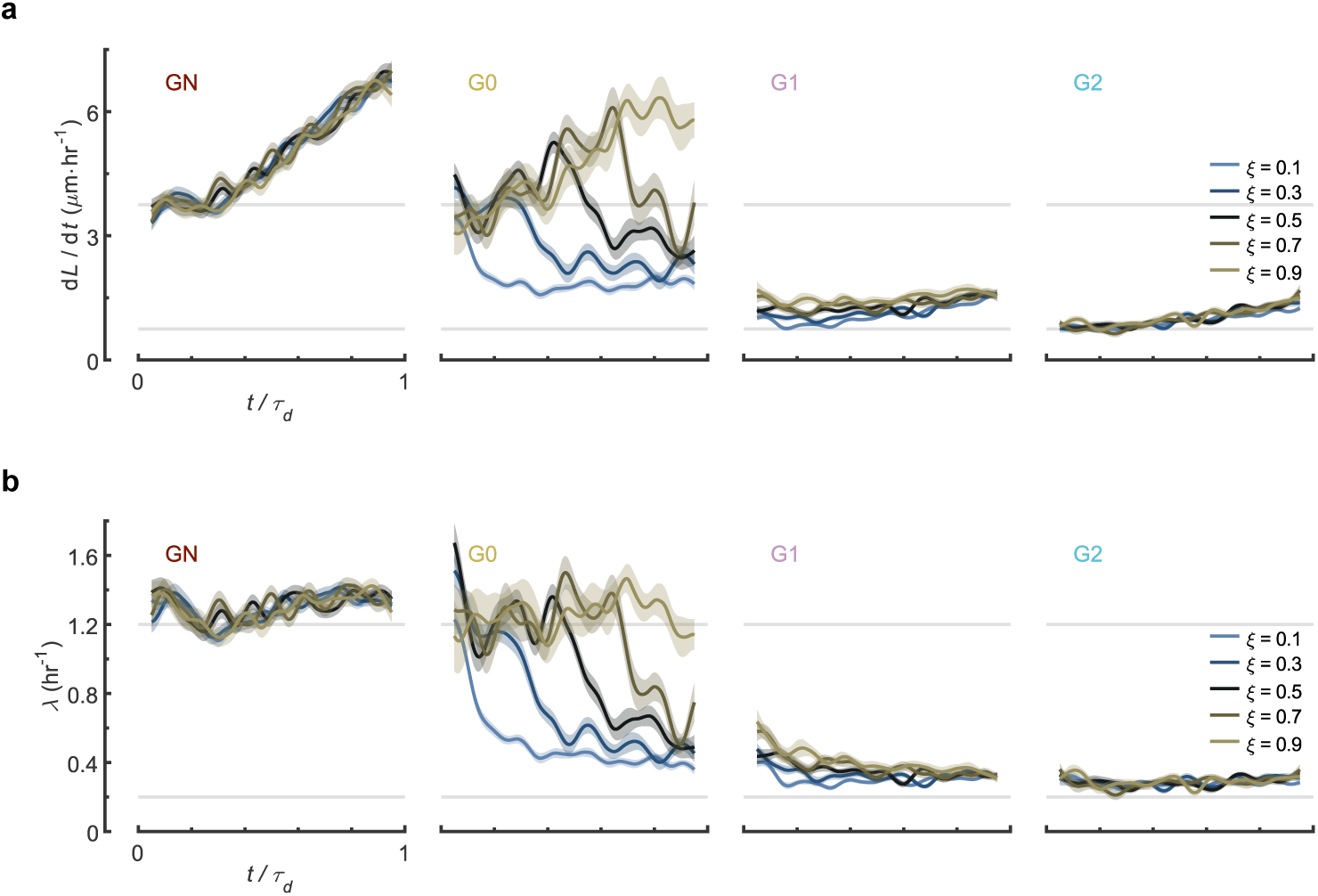
Dynamic details of grouped lineages. **a**, Elongation rate details of distinct *ξ* across generations. **b**, Growth rate details of distinct *ξ* across generations. In **a**, **b**, only 5 of 9 groups are shown; Error bands, s.e.m.

**Figure S13.**
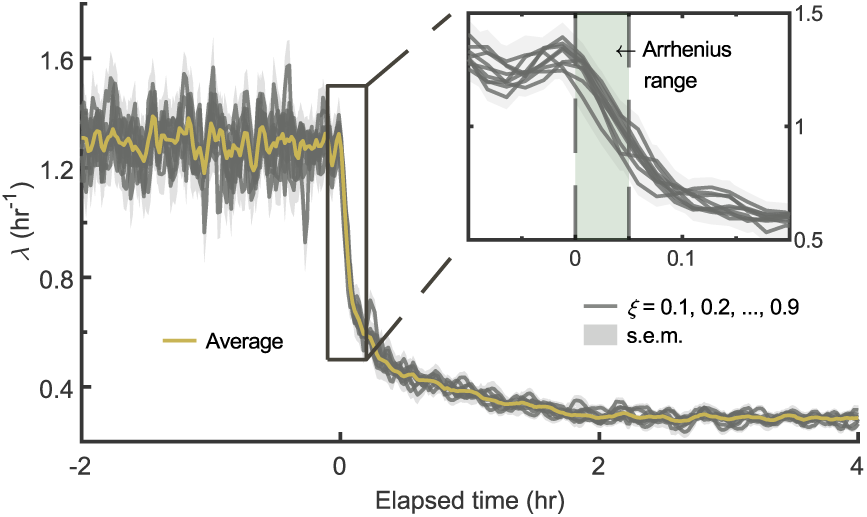
Growth dynamics of different group of cells. Average curve indicates the value of all cells. Arrhenius range of growth is shaded in inset. Error bands, s.e.m.

**Figure S14.**
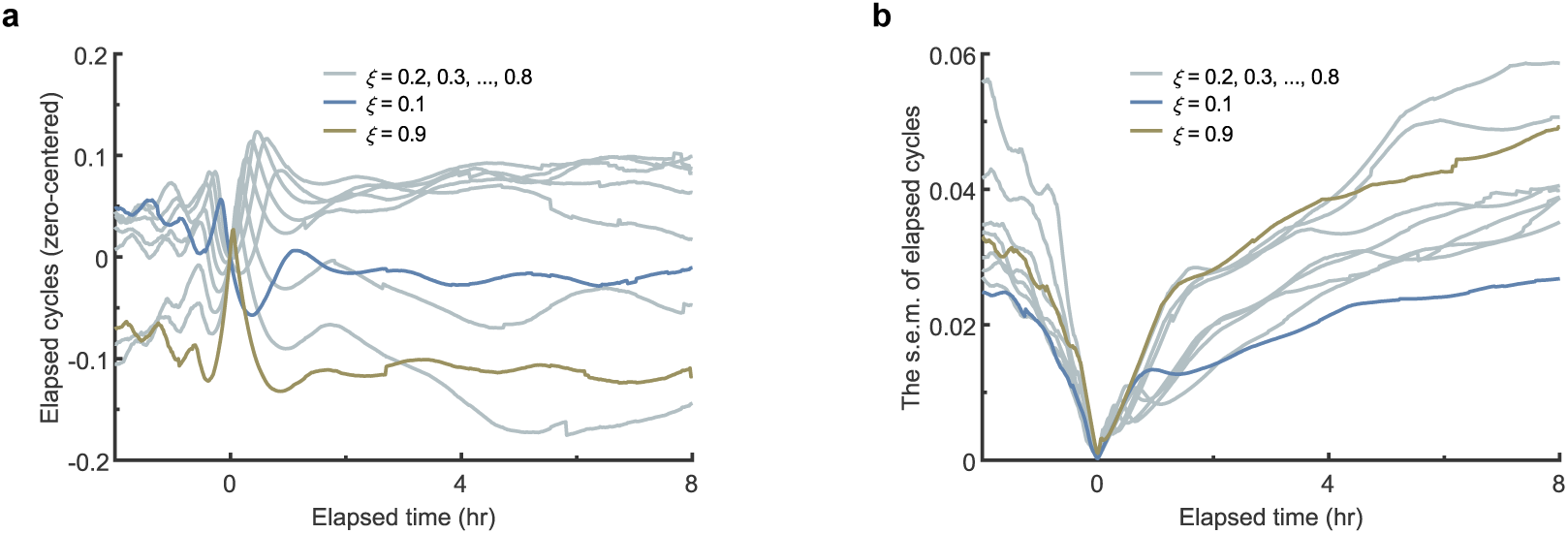
Synchronization in elapsed cycle number. **a**, Elapsed cycle numbers that centered by the average of all lineages as functions of time. **b**, Errors (s.e.m.) of elapsed cycle numbers as functions of time. Each group is centered by the average.

**Figure S15.**
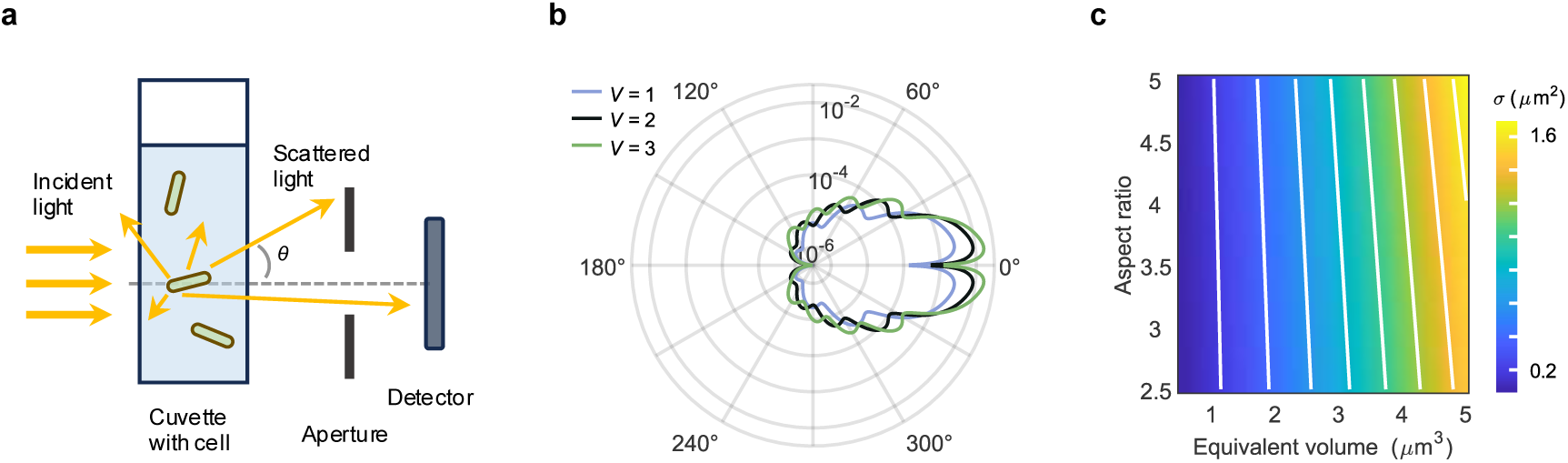
Numerical simulations based on scattering theory. **a**, Illustration of light scattering by a cell in the suspending medium. A small portion of scattered light is collected by the detector. **b**, Relative angular scattering intensity of single bacteria computed through the numerical simulation. *V*, equivalent volume in µm^3^. **c**, Simulation of cells with varying parameters. Color bar represents computed total cross section *σ* for each cell. White contour lines indicate the elevation of *σ*.

**Figure S16.**
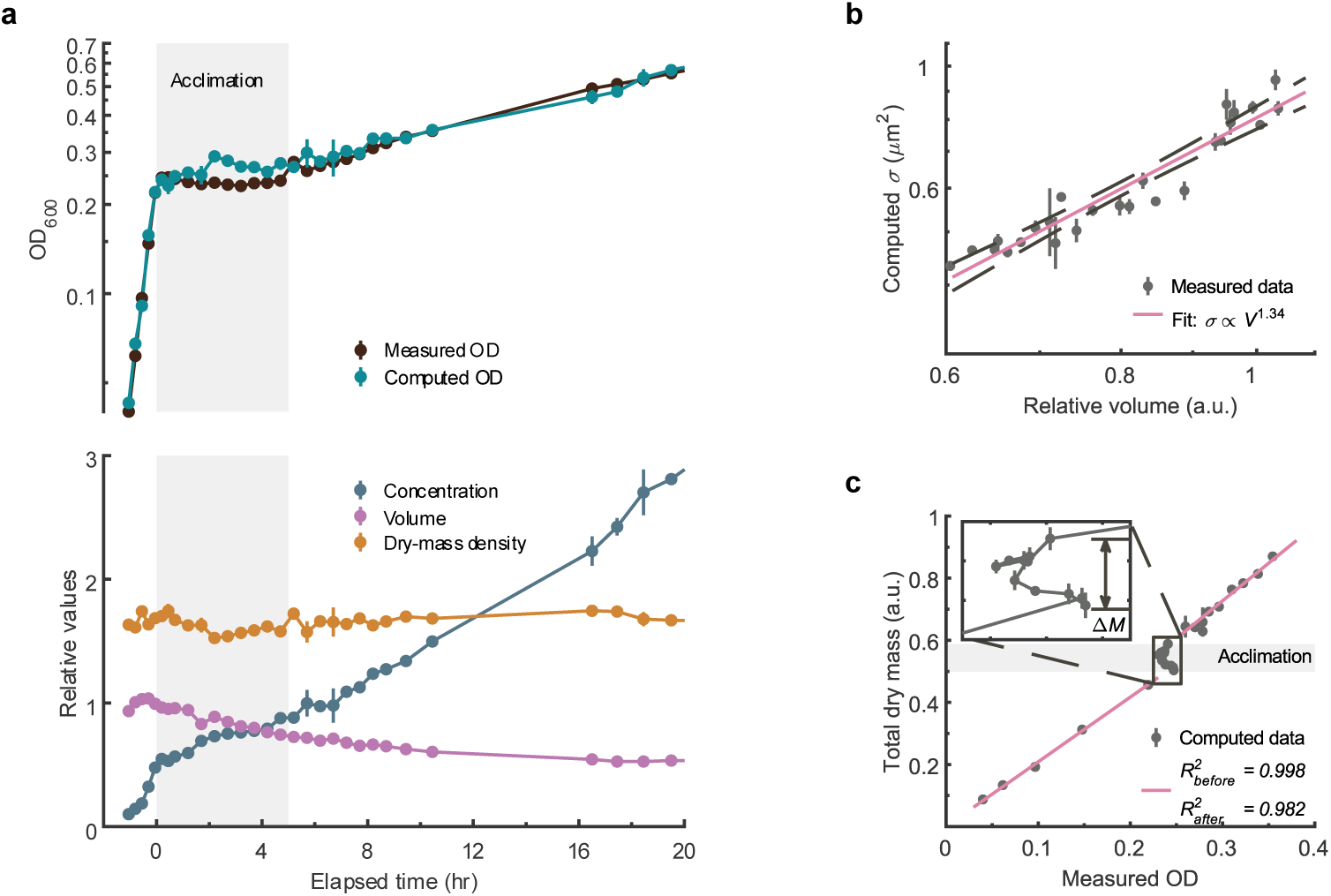
Supplemental results of batch culture. **a**, Measurements with temperature dropped to 12 ℃. Measured and computed OD_600_ values (top). Computed data was scaled at *t0*. Bottom: cell concentrations and average volumes extracted from the flow cytometer, normalized to *t0* and average value before CS, respectively. Relative cell density was computed and scaled for comparison. Shaded region indicates the acclimation. **b**, Computed *σ* from experimental data as function of relative volume. Power exponent was determined by linear regression on the logarithmically transformed data (*R^2^* = 0.913, confidence interval is ±0.17). Dashed lines indicate the 95% prediction bounds. a.u., arbitrary units. **c**, Computed total dry mass plotted as a function of OD, together with linear fittings. Fitting for the acclimation has a *R^2^* of -0.741 (Pearson correlation coefficient -0.59). Mass increment during the acclimation is indicated as the height of the shaded area and marked in the inset. Error bars in **a**-**c**, s.d.

**Figure S17.**
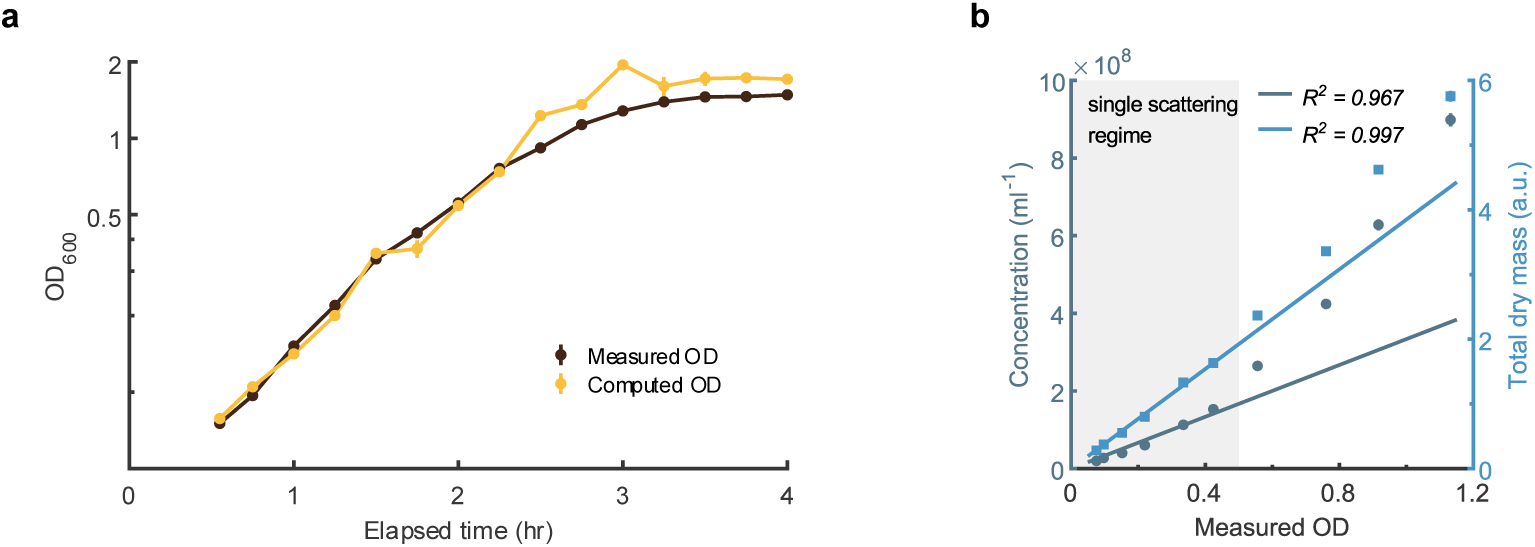
Supplemental results of batch culture without temperature downshift. **a**, Measured and computed OD_600_ in the batch culture without changing temperature. **b**, Correlation of measured OD_600_ in **a** with cell concentration and computed dry mass. Linear fittings were performed by data points within shaded area of which OD < 0.5. *R^2^* of the fitting is shown. Error bars in **a**, **b**, s.d.

**Figure S18.**
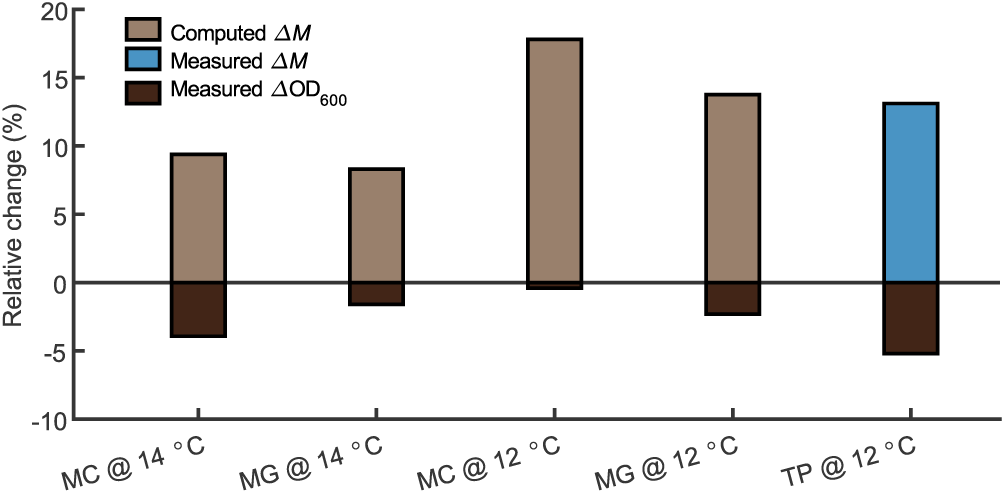
Percentage change of dry mass and OD_600_ during the acclimation. Temperature drops to 12 ℃ and 14 ℃ were performed separately for both strains: MC, MC4100 strain; MG, MG1655 strain. TP, total protein of MG1655 strain.

**Table S1.** LUNA performance results with different objective lenses from Nikon.

| Objective model | Plan Apo<br>$\lambda$ DM | Plan Apo<br>$\lambda$ DM | S Plan Fluor<br>ELWD | Super Fluor |
| --- | --- | --- | --- | --- |
| Immersion medium | oil | oil | air | air |
| Magnification (Mag) | 100× | 60× | 40× | 20× |
| Numerical Aperture (NA) | 1.45 | 1.40 | 0.60 | 0.75 |
| DOF ($\mu\text{m}$) | 0.46 | 0.54 | 1.80 | 1.41 |
| Focusing range ($\mu\text{m}$) | 18.6 | 40.1 | 111.5 | 400.8 |
| Linear range ($\mu\text{m}$) | 18.6 | 24.4 | 34.2 | 201.8 |
| Measurement repeatability (%) | 1.2 | 1.5 | 1.7 | 1.1 |
| Precision (nm) | 3.0 | 2.9 | 2.9 | 2.9 |
| Normalized precision (%) | 1.3 | 1.1 | 0.3 | 0.4 |

The DOF of the microscope was calculated using the formula $\text{DOF} = n \cdot \lambda / \text{NA}^2 + n \cdot s / \text{Mag}$ / NA. The refractive index $n = 1$ for the dry objectives, and $n = 1.515$ for the oil immersion objectives. Wavelength of illumination light $\lambda = 550$ nm. Pixel size of the imaging camera $s = 6.5$ $\mu\text{m}$ . Note that actual focusing range can be slightly larger than the values shown. Precision was calculated at a threshold set as 5 nm. Normalized precision was calculated as twice the ratio of Precision to DOF.

**Table S2.**
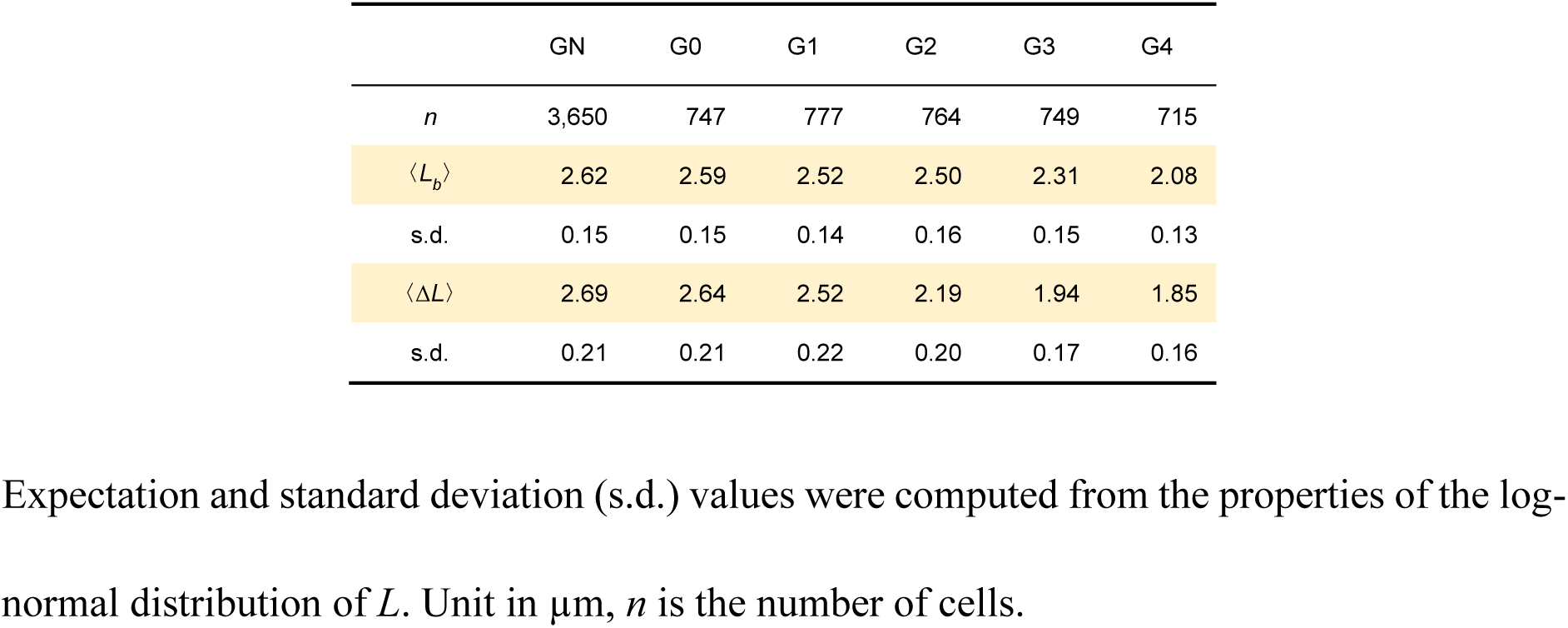
Length data of each generation.

## Video S1. Time-lapse imaging with slow and limited temperature downshift using commercial microscope

Image stack from the time-lapse experiment of *E. coli* cells with a commercial microscope (Nikon Instruments, Ti2 with autofocusing unit PFS). The experimental conditions are stated in the caption of Figure S1. The commercial autofocusing unit remain active throughout the experiment. Images were resized to 0.5 times.

## Video S2. LUNA-enabled single-cell level imaging with cold shock

Image stack from the time-lapse experiment of *E. coli* cells with LUNA-enabled bespoke microscope. The experimental conditions are stated in main text and methods. The stack is the same with shown in main text Figure. 2e. Images were resized to 0.5 times.

## Video S3. Montage of ten imaging sites of LUNA-enabled single-cell imaging with cold shock

The video presents ten montage microscopic fields of time-lapse imaging (randomly selected from the total 45 imaging sites). Over 200 single-cell lineages can be extracted to determine the cell viability in this video. Images were resized to 0.5 times.

